# Toxin-triggered activation of regulated exocytosis enhances bacterial egress from the intestinal layer

**DOI:** 10.64898/2026.02.06.704300

**Authors:** Christopher Margraf, Lilo Greune, Joyleen Fernandes, Paweena Wessel, Samriti Sharma, Johanna Sibbel, Sandra Heissler, Christian Rüter, Petra Dersch

## Abstract

Bacterial exit from host cells is essential for dissemination yet remains poorly understood. Here, we define a non-lytic egress pathway exploited by the enteric pathogen *Yersinia pseudotuberculosis* that co-opts the host exocytosis machinery in intestinal epithelial cells. The bacteria secrete a CNF-family toxin that activates the Cdc42-PLC γ1-IP_3_-IP_3_R signaling cascade, triggering SNARE-dependent fusion of the *Yersinia*-containing vacuoles with the cell membrane via VAMP7, Stx4, and SNAP23. This controlled exocytotic release preserves epithelial barrier integrity and occurs infrequently, representing a key rate-limiting step in systemic spread. These findings establish the host exocytotic machinery as an active determinant of bacterial egress, uncover a conserved vesicle trafficking pathway hijacked by intracellular pathogens for dissemination, and assign a new role for bacterial toxins in regulating host cell exit.

**Authors Summary:** Many bacterial pathogens penetrate and cross protective host cell barriers to spread within the body. While the mechanisms by which enteric bacteria enter host cells are well characterized, far less is known about how they exit host cells after invasion. In this study, we investigated how the enteric pathogen *Yersinia pseudotuberculosis* exits from human gut epithelial cells after crossing them. We found that they egress at the basolateral side of the intestinal cells through a controlled, non-destructive process that preserves the integrity of the epithelial barrier. To achieve this, the bacteria co-opt host signaling pathways involved in host cell vesicle release (exocytosis) to promote controlled egress. These exit events are rare, suggesting that bacterial escape from cells is a major bottleneck during infection. Importantly, we show that this process is enhanced by a secreted toxin of the Cytotoxic Necrotizing Factor (CNF) family (CNF_Y_), which promotes the fusion of the bacteria-containing vacuole with the basolateral cell membrane. This finding uncovers a previously unrecognized role of bacterial toxins in facilitating bacterial cell egress.

## Introduction

Many microbial pathogens invade and manipulate the intracellular environment of host cells to promote intracellular survival and proliferation, evade host immune responses, or breach barriers to disseminate to adjacent tissues. Over the past few years, considerable progress has been made in unraveling the versatile and complex strategies of pathogens to invade their host cells and establish an intracellular niche (e.g., within a vacuole/phagosome or the cytoplasm). In contrast, our knowledge about how pathogens egress from their host cells is still in its infancy, due to experimental hurdles (e.g., difficult genetic manipulations of differentiated host cell types and pathogens) and the perception of this process as passive. For instance, in *Legionella pneumophila,* it was initially assumed that the metabolic and mechanical burden imposed by the massive number of intracellular bacteria could lead to cell lysis and the passive release of the pathogen [1].

However, recent studies demonstrated that host cell exit by pathogens is an active and highly dynamic process promoted by multiple pathways involving diverse molecular mechanisms used by bacteria, fungi, and protozoa (reviewed in [2–6]). To date, the following major exit strategies have been described: (i) the induction of programmed cell death, e.g., pyroptosis, necroptosis, apoptosis, (ii) active host cell destruction, e.g., host cell lysis triggered by microbial pore-forming toxins or proteases, and (iii) membrane-dependent exit processes without cell lysis, including e.g., exocytosis, expulsion, extrusion, budding, and actin-mediated protrusion. A substantial number of bacteria have been found to activate cell death programs to exit from host cells, including *Mycobacterium, Burkholderia, Francisella, Legionella, Listeria, Salmonella,* and *Shigella* species [3,4]. The high prevalence of specific exit pathways suggests a convergent evolution of strategies that appear to confer a selective advantage to the microbe [4]. One species can also trigger distinct exit pathways in different host cell types or different infection stages [3,4]. For instance, *Listeria monocytogenes* can trigger pyroptosis upon infrequent bacteriolysis in the host cell cytosol [7]. They can exit cells by forming actin tails, which they use to project from the infected cell within the tip of a filopodium-like membrane extension. This protrusion is engulfed by the adjacent cell without contact of the bacteria with the extracellular environment [8].

There is compelling evidence that enteric human pathogenic *Yersinia* species, such as *Y. entercolitica* and *Y. pseudotuberculosis,* enter and transfer through specialized epithelial microfold (M) cells that reside within the follicular-associated epithelium above Peyer’s patches of the intestinal epithelial layer to gain access to underlying lymphatic tissues [9–12]. M cells are specialized intestinal epithelial cells that play a crucial role in immunosurveillance [13]. They ingest luminal contents/antigens and deliver them to immune cells that reside closely to their cell body due to the invaginated basolateral side [10]. In contrast to the adjacent enterocytes, they lack apical microvilli and express β_1_-integrins on their apical surface [10–12]. Several studies have shown that enteropathogenic *Yersiniae* bind to β_1_-integrins and enter M cells primarily via the bacterial surface adhesin invasin (InvA), and other *Yersinia* adhesins can support this process [9,12,14,15]. However, our understanding of how bacteria reside, survive, and exit M cells is limited due to the lack of suitable cell culture systems. Very recently, a study utilized human ileal enteroid-derived monolayers containing five different intestinal cell types, including M cells, to investigate the interaction between *Y. pseudotuberculosis* and M cells [14]. They found that *Y. pseudotuberculosis* specifically targets M cells, recapitulating murine studies. Interestingly, transcytosis was significantly repressed by host-adapted *Y. pseudotuberculosis* expressing the *Yersinia* outer membrane proteins (Yops) and the Ysc-type III secretion system (Ysc-T3SS) with which they are injected into host cells [14,16,17].

Reduction of transcytosis occurred due to YopE and YopH preventing the uptake of *Y. pseudotuberculosis*, as well as YopE-mediated extrusion of M cells from the epithelium [14]. This finding suggested that efficient *Y. pseudotuberculosis* transit occurs before high levels of Ysc-T3SS/Yop expression are reached.

Despite our increasing understanding of *Yersinia*-M cell interactions, triggered invasion and transcytosis, the molecular mechanism of bacterial exit from these cells on the basolateral side remains poorly understood. In this study, we used human intestinal monolayers containing M cells to reconstruct the intestinal monolayer and visualize how *Y. pseudotuberculosis* crosses it to gain access to the underlying environment. We show that the bacteria egress via exocytosis without inducing cell death or disrupting monolayer integrity. The efficiency of transcytosis is significantly enhanced by the CNF_Y_ toxin, which initiates the Ca^2+^-regulated exocytotic process by activating the PLC1-IP3-IP3R signaling cascade through the Rho GTPase Cdc42. This promotes the interaction of bacteria-containing vesicles (bacterial phagosomes) with the plasma membrane via the vesicle SNARE (v-SNARE) protein VAMP7, the target SNARE (t-SNARE) protein Stx4, and the synaptosomal-associated protein SNAP23, which are essential for exocytosis. In summary, this study provides visual, mechanistic, and functional insights into the egress of *Y. pseudotuberculosis* from M cells in the human intestinal epithelium. It demonstrates that the clonal invasion and abscess formation in the Peyer’s Patches is likely initiated by rare cell exit events and not solely by an efficient initial immune response to *Yersinia* infection, as previously thought.

## Results

### *Y. pseudotuberculosis* passes rapidly through intestinal epithelial cells via non-lytic bacterial egress in a T3SS/Yop-independent manner

The success of *Y. pseudotuberculosis* as an enteropathogen is not only linked to its invasion of intestinal epithelial cells at the apical side; it also depends strongly on its transcytosis and egress from these cells at the basolateral side to reach underlying and deeper tissues. To elucidate the molecular mechanisms involved in *Yersinia* epithelial cell exit, we established a Caco-2 cell-based monolayer, including M cells, in a transwell system similar to that described [18–20] (**Fig. 1A**), which has previously been used to study particle uptake and bacterial transcytosis [21,22]. The Caco-2 monolayer was characterized by the presence of microvilli on the apical side, the formation of distinct cell-cell contacts, and dense basolateral adherence to the transwell membrane. The presence of cells displaying morphological and functional characteristics of M cells in the monolayer was confirmed by confocal microscopy testing for M cell markers (e.g., Glycoprotein-2/GP2), and they were predominantly recognized and bound by *Y. pseudotuberculosis* (**Fig. 1B-C**). The intactness/integrity of the monolayer was ensured by (i) the use of 40 kDa TRITC-Dextran to visualize that particle diffusion through the intestinal cell barrier is prevented (Fig. **S1A**) and (ii) a continuous measurement of the transepithelial electrical resistance (TEER) (Fig. **S1B**).

**Figure 1:**
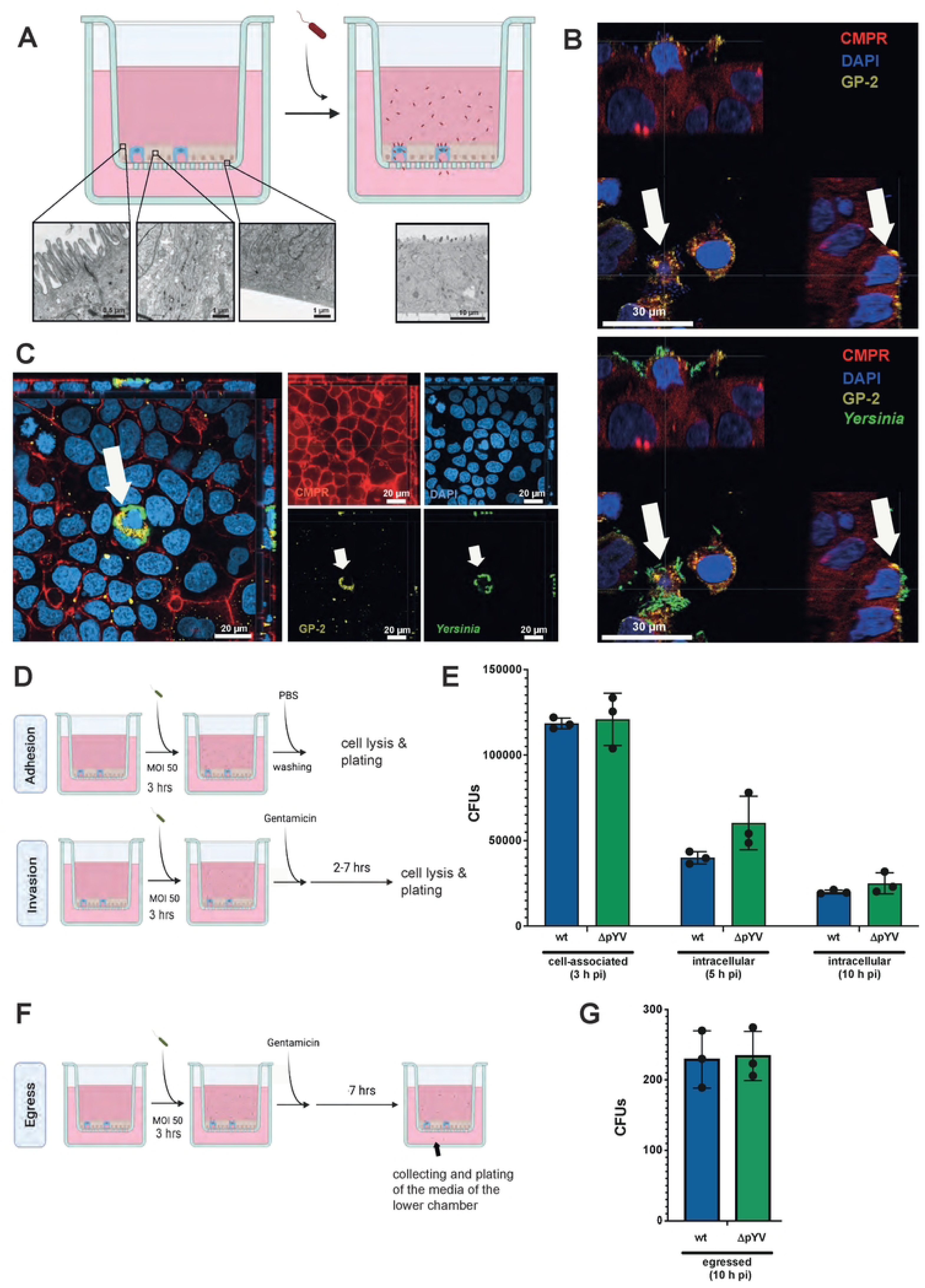
Establishment of a standardized human intestinal epithelial cell monolayer system for the analysis of bacterial transcytosis. (**A**) Illustration of the Caco-2 2D cell monolayer transwell system. Caco-2 cells were seeded into transwell inserts (3 µm pore size) and cultivated until the TEER had reached > 300 Ω•cm². Subsequently, the monolayer was infected with the *Y. pseudotuberculosis* wildtype strain YPIII at an MOI of 50. Differentiation (e.g., microvilli formation; left panel below), monolayer formation with adherent bacteria (right panel below), and barrier integrity (both middle panels) were visualized by electron microscopy. Created in BioRender. Dersch, **P.** (2026). (**B,C**) Confocal laser scanning microscopy (C-LSM) of Caco-2 monolayer infected with *Y. pseudotuberculosis* strain YPIII pFU92 expressing GFP (green) and stained with DAPI (nuclei-blue), and CellMask Plasma Membrane Red (CMPR) (red), and antibodies against glycoprotein-2 (GP-2) (yellow). Orthogonal XZ and YZ planes (**B**) and the apical surface of the infected Caco-2 monolayer (**C**) are shown. (**D**) Illustration of the methods used to determine the adherent/cell-associated number and intracellular number of *Y. pseudotuberculosis* in the Caco-2 cell monolayer after infection. The monolayer was infected with 2.5 x 10^6^ bacteria of *Y. pseudotuberculosis* wildtype strain YPIII (wt) or the virulence plasmid-negative strain YP12 (ΔpYV) (MOI of 50), incubated for 3 h, and (i) washed with PBS and lysed to determine the number of cell-associated bacteria post-infection (pi), or (ii) gentamicin was added for an additional 2-7 h (i.e., 5 h and 10 h post-infection) before the cells were lysed, and the number of cell-associated or intracellular bacteria was quantified by plating serial dilutions on LB plates. Created in BioRender. Dersch, **P.** (2026). (**E**) The CFUs of the cell-associated and intracellular bacteria at the indicated time points are given. (**F**) Illustration of the methods used to determine the number of *Y. pseudotuberculosis* that egressed on the basolateral side of the Caco-2 cell monolayer. (**G**) 7 h after gentamicin was added to the infected monolayer (i.e., 10 h post-infection), the medium containing the egressed bacteria was collected and plated on LB plates. Created in BioRender. Dersch, **P.** (2026). (**E,G**) The mean +/- SEM of three independent biological replicates is shown. Significant differences were determined using a two-way ANOVA test with Tukey correction.

To quantify and characterize the egress of *Y. pseudotuberculosis*, differentiated Caco-2 monolayers were infected apically with the wildtype (wt) strain YPIII at a multiplicity of infection (MOI) of 50 (2.5 x 10^6^ bacteria per layer of 5 x 10^4^ cells). To mimic initial host cell contact, we pre-grew the bacteria at a moderate temperature at which the bacteria expressed the primary adhesion and invasion factor invasin (InvA) [23]. We added them to the monolayer without centrifugation to allow loose fimbria-mediated cell contact, avoid mechanical cell stress/damage, and avoid artificial induction of the T3SS/Yop machinery. After 3 h, gentamicin was added to the apical medium to stop the ongoing invasion and reinfection of apically egressed bacteria (**Fig. 1D**).

To assess the efficiency and dynamics of this process, we first determined the number of associated and intracellular bacteria at 3 h, 5 h, and 10 h post-infection by plating the infected cell lysates (**Fig. 1D-E**). We found that a suitable number of bacteria (approximately 4-5% of added bacteria) had bound and invaded the epithelial cells 3 h post-infection (**Fig. 1E**), particularly those with M cell characteristics (**Fig. 1B-C**). Moreover, we observed substantially reduced intracellular bacteria over time (1-2% is intracellular 10 h post-infection, **Fig. 1E**). Bacterial-mediated lysis of the intestinal cells and destruction of the monolayer were ruled out by continuous measurements of the TEER (Fig. **S1B**) and lactose dehydrogenase release/cell toxicity assays (**S1C**), which indicated no monolayer damage within 10 h post-infection.

To test whether we can identify bacteria that have egressed basolaterally, we initially collected the basal medium 10 h post-infection. We determined the number of bacteria (colony-forming units/CFUs) by plating (**Fig. 1F**). Approximately 200-300 bacteria were detected in the basolateral chamber (**Fig. 1G**). Although the proliferation of bacteria in the cell culture medium is relatively slow (doubling time of > 3 hours), we aimed to exclude this variable to ensure accurate quantification of basolaterally egressed bacteria over time. For this purpose, we collected the basal medium every 30 minutes post-infection to plate and quantify egressed bacteria (colony-forming units/CFUs), and replaced the basal chamber with fresh antibiotic-free medium. This collection procedure was repeated every 30 minutes for 10 hours to establish a temporal profile of *Y. pseudotuberculosis* egress from the Caco-2 cells. As shown in **Fig. 2A**, the bacteria began to egress approximately 2 hours after infection, with a linear rise to a strong release peak at 3.5 hours post-infection, followed by a linear decline until 5 hours post-infection. From the initial cell-bound bacteria (10^5^ CFUs), 200-300 egressed on the basolateral side (approximately 0.02%). The exit pattern did not change when gentamicin was added at different time points (e.g., 2, 3, or 4 hours post-infection, **Fig. 2A**), indicating that the antibiotic treatment does not influence the release peak. Based on these results, we determined the overall number of bacteria that had egressed, generally 5 hours post-infection, in the following experiments.

**Figure 2:**
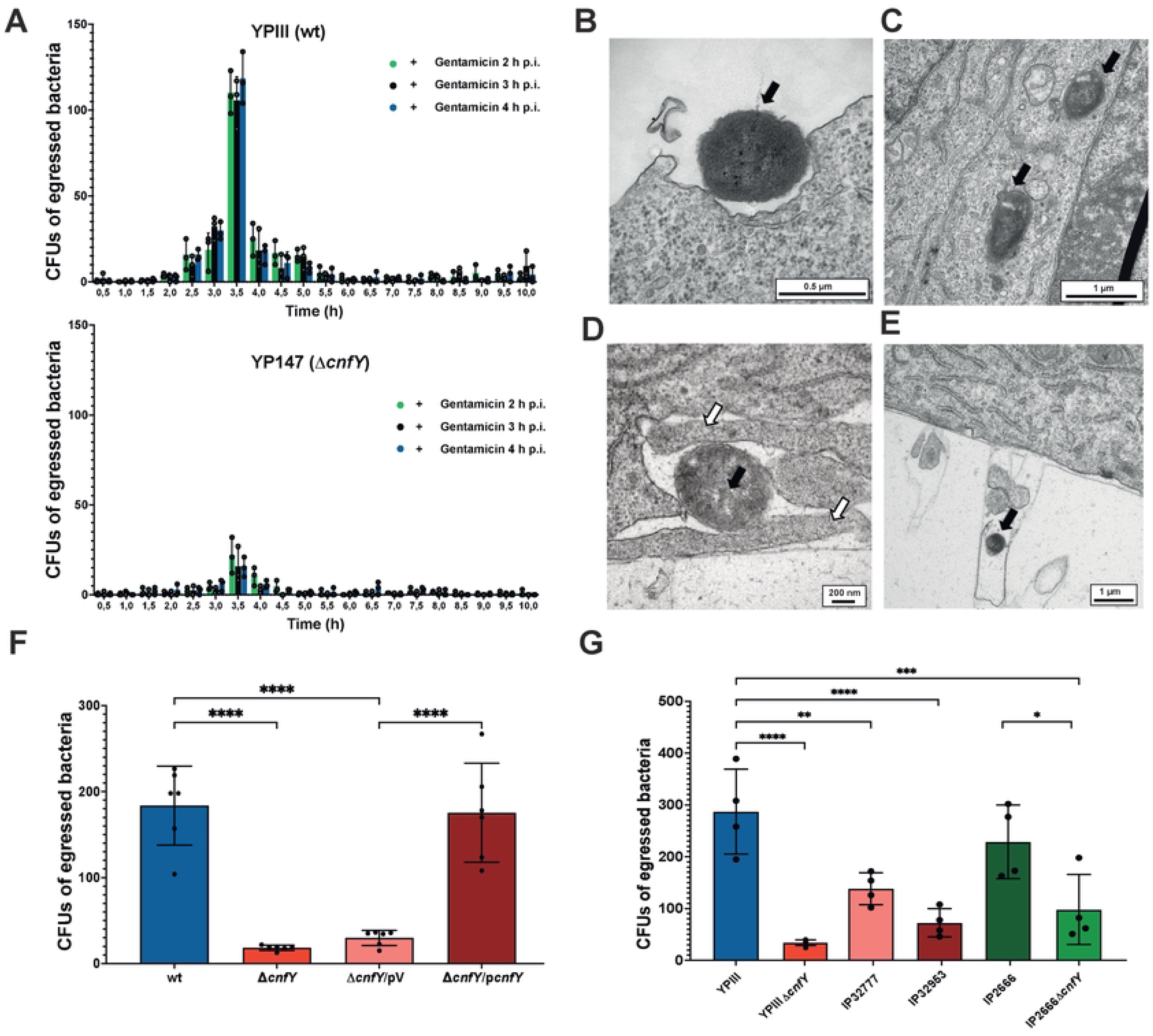
Translocation of *Y. pseudotuberculosis* across the Caco-2 monolayer peaks at 3.5 h post-infection, and depends on the CNF_Y_ toxin. (**A**) Caco-2 2D monolayer was infected with 2.5 x 10^6^ bacteria of the *Y. pseudotuberculosis* wildtype strain YPIII (wt) (upper panel) or the YP147 (Δ*cnfY*) mutant strain (lower panel). After 2, 3, or 4 h post-infection (p.i.), gentamicin was added to kill extracellular bacteria. Every 30 minutes post-infection, the medium from the basolateral chamber was collected over 10 hours and replaced. The removed medium was then plated on LB plates to determine the CFUs of egressed bacteria at the indicated time point. The mean +/- SEM of three independent experiments is shown. (**B-E**) Caco-2 2D monolayer was infected with *Y. pseudotuberculosis* wildtype strain YPIII, and monolayer samples were taken after 3.5 h and analyzed by transmission electron microscopy. Cell-adherent bacteria (**B**), as well as intracellular *Yersiniae* (**C**) were identified, indicated by black arrows. (**D**) *Yersinia* during the exit process at the basolateral side of the Caco-2 cells or (**E**) *Yersinia* that has successfully egressed from the Caco-2 monolayer are shown; white arrows indicate lamellipodia often associated with the egressing bacteria. (**F**) Caco-2 monolayer was infected with the *Y. pseudotuberculosis* wildtype strain YPIII (wt), the YP147 (Δ*cnfY*) mutant strain +/- the empty vector pJNS11 (pV), or the *cnfY*^+^ complementation plasmid pJNS10 (p*cnfY*). 3 hours post-infection, gentamicin was added to kill extracellular bacteria. After an additional 30 minutes of incubation, the medium from the basolateral chamber was collected and plated on LB plates to determine the CFUs of egressed bacteria. The mean +/- SEM of six independent biological replicates is shown. Significant differences were determined using a two-way ANOVA test with Šidák correction and indicated by asterisks (P-value: **** < 0.0001). (**G**) Comparative study of bacterial egress of strains with and without a functional *cnfY* toxin gene. Caco-2 monolayer was infected with 2.5 x 10^6^ bacteria of *Y. pseudotuberculosis* wildtype strain YPIII (*cnfY*^+^), YP147 (YPIII Δ*cnfY*) mutant strain, IP32777 (*cnfY*^-^), IP32953 (*cnfY*^-^), IP2666 (*cnfY*^+^), and the IP2666 Δ*cnfY* mutant strain. 3 hours post-infection, gentamicin was added to kill extracellular bacteria. After an additional 30 minutes of incubation, the medium from the basolateral chamber was collected and plated onto LB agar to determine the CFUs of egressed bacteria. The mean +/- SEM of four independent biological replicates is shown. Significant differences were determined using a two-way ANOVA test with Tukey correction and indicated by asterisks (P-value: * < 0.05, ** < 0.01, *** < 0.001, **** < 0.0001).

To gain further insight into the fate of the bacteria within host cells and the mechanism of egress, we determined their cellular localization and the egress process using electron microscopy. We found adherent bacteria on the apical side of the Caco-2 cells, and invading and intracellular bacteria localized in membrane-bound vacuoles during their passage across the cells (**Fig. 2B-C**). We further identified bacteria at the site of egress on the basolateral side of human epithelial cells, frequently associated with membrane protrusions resembling filopodia and lamellipodia-like structures (**Fig. 2D**). Additionally, we observed egressed bacteria on the basolateral side of the monolayer (**Fig. 2E**). Moreover, we found that many intracellular bacteria are lysed during transcytosis (Fig. **S2A**), and severely infected cells were expelled via epithelial extrusion, as previously described [15][24] (Fig. **S2B**). This explains the observed reduction of viable intracellular bacteria over time.

To identify bacterial factors that are implicated in the translocation of the epithelial layer, we started to test the role of the virulence factors encoded by the *Yersinia* virulence plasmid pYV, encoding the Ysc-T3SS/Yop effectors known to manipulate cell responses [14,16,17] by comparing *Yersinia* adhesion, invasion and egress of the pYV-cured strain (ΔpYV) with that of the wildtype (wt). Over the entire tested infection period, we detected no cytotoxic effects from the infection and antibiotic treatment (Fig. **S1C**). Moreover, we found that adhesion, invasion/intracellular persistence, as well as translocation/egress were comparable between the wildtype and the pYV-cured strain (**Fig. 1E,G**). This finding indicated that the virulence plasmid-encoded T3SS/Yop factors are dispensable for *Yersinia* transcytosis.

### The *Yersinia* toxin CNF_Y_ promotes translocation across the intestinal epithelial cell monolayer

Having shown that the *Yersinia* virulence plasmid does not influence transcytosis, we examined whether other virulence-relevant factors of *Yersinia* play a significant role in this process. Using our established setup, we tested our collection of *Y. pseudotuberculosis* mutants deficient in individual virulence-relevant factors. We found that a mutant strain deficient in the cytotoxic-necrotizing factor of *Yersinia* CNF_Y_ (Δ*cnfY*) was significantly reduced in its ability to translocate through the Caco-2 cell monolayer (**Fig. 2F**). Complementation of the gene defect by the integration of a *cnfY*^+^-encoding plasmid, but not the empty vector (pV), regenerated the efficiency of translocation to nearly wildtype levels (**Fig. 2F**). Loss of this toxin did not result in lower numbers of cell-bound and intracellular bacteria (Fig. **S3A**) and did not compromise the host cell viability throughout the experiment (Fig. **S3B**). On the contrary, a slightly higher number of intracellular bacteria was identified within the Caco-2 monolayer (Fig. **S3A**), as expected when the facilitating influence of the toxin on bacterial egress is absent. A time course of the Δ*cnfY* mutant egress over 10 h revealed a release pattern comparable to that of the WT, although overall, much fewer bacteria were released (**Fig. 2A**). This analysis showed that the overall egress, rather than the exit dynamics, changed after CNF_Y_ treatment, indicating that the CNF_Y_ toxin is not essential for transcytosis but strongly enhances this process.

It has previously been shown that specific *Y. pseudotuberculosis* isolates, such as YPIII and IP2666, encode the entire *cnfY* gene, but many isolates also harbor deletions that eliminate toxin function [25]. If CNF_Y_ indeed promotes epithelial transcytosis, *cnfY*-deficient isolates should exhibit a reduced ability to translocate across the intestinal epithelial layer. To test this hypothesis, we compared the ability to transcytose and egress from the Caco-2 monolayer of two clinical *Y. pseudotuberculosis* isolates (IP32777, IP32953) that lack a functional *cnfY* gene with two strains (YPIII and IP2666) that possess a functional *cnfY* copy and their isogenic *cnfY* mutants. In fact, the number of egressed bacteria was significantly higher in CNF_Y_-positive strains than in CNF_Y_-negative strains or *cnfY* mutant variants (**Fig. 2G**), whereas bacterial adherence and epithelial cell integrity were comparable among all strains (Fig. **S4**). Moreover, the intracellular bacterial load was also higher in the CNF_Y_-negative strains (Fig. **S4**), similar to the Δ*cnfY* mutant variants of YPIII and IP2666 (Fig. **S3, S4**),

### CNF_Y_-promoted *Yersinia* egress via activation of the Rho GTPase Cdc42

The CNF_Y_ toxin is a single-chain A-B toxin comprising an N-terminal delivery domain and a C-terminal activity domain. Upon endosomal uptake into the host cell endosome, the toxin undergoes autocatalytic cleavage, releasing the active domain into the host cell cytosol [25,26]. CNF_Y_ deamidates Gln61/63 of the small Rho GTPases RhoA, Rac1, and Cdc42, resulting in constitutively activated Rho GTPases with a blocked GTP hydrolase activity [27–29]. The constitutively activated deamidated Rho GTPases induce actin polymerization, leading to the formation of membrane protrusions, including lamellipodia, filopodia, and stress fibers [25,28]. As substantial evidence exists that Rho GTPases are implicated in the remodelling of actin as well as docking and fusion of vesicles with the plasma membrane during (polarized, regulated) exocytosis [30–32], we addressed the role of the individual Rho GTPases in *Yersinia* transcytosis and egress. For this purpose, we infected the Caco-2 monolayer with *Y. pseudotuberculosis* (wt), the CNF_Y_-deficient (Δ*cnfY*) strain, the complemented strain (Δ*cnfY*/p*cnfY*), and the empty vector control strain (Δ*cnfY*/pV). After three hours, the medium in the apical chamber was replaced with a gentamicin-containing medium supplemented with specific inhibitors targeting RhoA or Cdc42/Rac1. The delayed addition of the Rho GTPase inhibitors ensured that bacterial adhesion and entry remained unaffected, as it has been previously demonstrated that Rho GTPases, particularly Rac1, regulate this process [33]. Inhibition of RhoA did not decrease *Yersinia* transcytosis and egress (Fig. **S5A**).

In contrast, inhibition of Cdc42/Rac1 resulted in a significant and substantial reduction of this process in both the wild-type and the Δ*cnfY*/p*cnfY* complemented strain (**Fig. 3A**), while the number of cell-associated bacteria remained unaffected (Fig. **S6A**). Similar to the loss of CNF_Y_, inhibition of Cdc42/Rac1 activation resulted in a slight but significant increase in intracellular bacteria (Fig. **S6A**), likely due to reduced bacterial egress. The addition of the Cdc/42/Rac1 inhibitor did not further reduce the low but still detectable levels of Δ*cnfY* egress, indicating that CNF_Y_-mediated activation of *Yersinia* transcytosis and/or egress is mainly driven by its effect on Rho GTPases.

**Figure 3:**
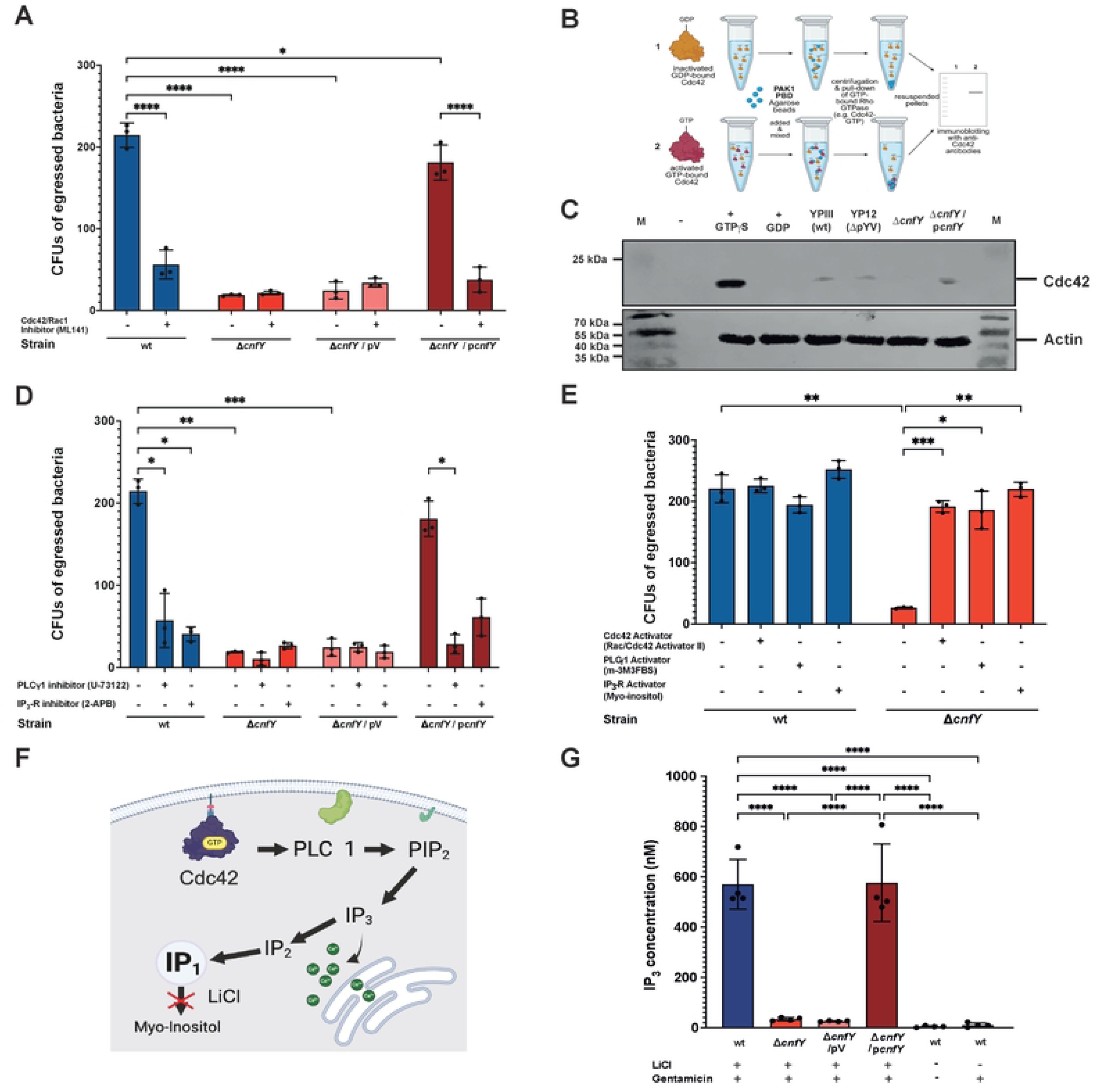
CNF_Y_-mediated activation of Rho GTPase Cdc42, PLC γ-1, and IP3-R enhances bacterial egress. Caco-2 monolayer was infected with 2.5 x 10^6^ bacteria of the *Y. pseudotuberculosis* wildtype strain YPIII (wt), the YP147 (YPIII Δ*cnfY*) +/- the empty vector pJNS11 (pV), and the *cnfY*^+^ complementation plasmid pJNS10 (p*cnfY*). (**A**,**D**,**E**) 3 hours post-infection the medium was removed and replaced by medium containing gentamicin and inhibitors of the Cdc42/Rac1 (ML141) (**A**), phospholipase C (PLC γ-1) (U-73122) (**D**), and Ca^2+^ channel IP3-R (2-APB) (**D**), or activators for Cdc42/Rac1 (Activator II), PLC γ-1 (m-3M3FBS), or IP3-R (Myo-Inositol) (**E**). After 30 minutes, the medium from the basolateral chamber was collected and plated onto LB agar to determine the CFUs of egressed bacteria. The mean +/- SEM of three independent biological replicates is shown. Significant differences were determined using a two-way ANOVA test with Tukey correction and indicated by asterisks (P-value: * < 0.05, ** < 0.01, *** < 0.001, **** < 0.0001). (**B**,**C**) Caco-2 monolayer was infected with 2.5 x 10^6^ bacteria of the *Y. pseudotuberculosis* wildtype strain YPIII (wt), the *Yersinia* virulence-negative strain (ΔpYV), the *cnfY*-negative mutant strain YP147 (Δ*cnfY*) without or with *cnfY*^+^ complementation plasmid pJNS10 (p*cnfY*). After 3.5 h post-infection, infected Caco-2 cells were lysed, cell extracts were prepared. Activation of the different Rho GTPases was tested by the isolation of the GTP-bound form by pull-downs with Rho-GTPase-binding agarose beads and Western blotting using Rho GTPase-specific antibodies, e.g., against Cdc42. M: Protein size marker. As negative and positive controls, high concentrations of GDP and GTP (GTP γS) were added to the uninfected whole-cell extract samples. Equal concentrations of extracts were used for pull-down assays, which were assessed by Western blotting with Actin antibodies. (**B**) shows a scheme of the procedure (Created in BioRender. Dersch, **P.** (2026)), and (**C**) the Western blots; (**F**,**G**) CNF_Y_-mediated induction of inositol triphosphate (IP_3_) production. (**F**) Scheme of triggered Cdc42-mediated activation of phospholipase C (PLC γ-1), which leads to the formation of inositol monophosphate (IP_3_) from PIP_2._ IP_3_ is rapidly metabolized to inositol monophosphate (IP_1_), and IP_1_ can thus be used as a proxy for IP_3_ levels by adding LiCl, which blocks the metabolism of IP_1_. Created in BioRender. Dersch, **P.** (2026). (**G**) Caco-2 monolayer was infected with 2.5 x 10^6^ bacteria of the *Y. pseudotuberculosis* wildtype strain YPIII (wt), YP147 (YPIII Δ*cnfY*) +/- the empty vector pJNS11 (pV), and the *cnfY*^+^ complementation plasmid pJNS10 (p*cnfY*). After at least 2 h post-infection, the medium was removed and replaced with medium +/- gentamicin and/or 50 mM LiCl to block IP_1_ metabolism, as indicated. After an additional 1.5 h, the Caco-2 cells were lysed, and the cellular concentration of IP_1_ was determined by ELISA using an anti-IP_1_ monoclonal antibody. The IP_3_ concentrations were calculated based on the IP_1_ amounts. The mean +/- SEM of four independent biological replicates is shown. Significant differences were determined using a two-way ANOVA test with Tukey correction and indicated by asterisks (P-value: **** < 0.0001).

Next, we examined whether Cdc42 or Rac1 is responsible for the observed activation of *Yersinia* translocation. For this purpose, we harvested the Caco-2 cells 3.5 hours after infection with the WT, the Δ*cnfY* mutant, and the complemented strain Δ*cnfY*/p*cnfY.* We examined the amount of activated (GTP-bound) RhoA, Cdc42, and Rac1 by Western blot analysis (**Fig. 3B**). No activation of RhoA (Fig. **S5B**) and only a very weak activation of Rac1 could be detected upon infection with the complemented CNF_Y_-overexpression strain Δ*cnfY*/p*cnfY* (Fig. **S5C**). In contrast, GTP-bound active Cdc42 was detectable in lysates of WT and Δ*cnfY*/p*cnfY-*infected cells but not in lysates of cells infected with the Δ*cnfY* mutant strain (**Fig. 3C**). Together, this indicated that CNF_Y_-triggered activation of *Yersinia* translocation depends mainly on the activation of the Rho GTPase Cdc42.

### CNF_Y_ activates *Yersinia* egress via Cdc42-triggered PLCγ1 and IP_3_-receptor activation

Several studies addressing the role of Rho GTPases in the context of exocytotic processes have elucidated that RhoA, Rac1, and Cdc42 localize to intracellular secretory vesicles and the secretion machinery, the exocyst [31,34]. The exocyst is a multiprotein complex that functions in the precise targeting, docking, and fusion of vesicles from endosomal compartments with the plasma membrane to release their cargo, a process called polarized exocytosis [32]. Polarized exocytosis requires well-coordinated actomyosin dynamics to efficiently transport vesicles by the trafficking machinery to the release site. Rho GTPases act in concert to accomplish this task [32,34]. Based on this knowledge, we hypothesized that the CNF_Y_ toxin of *Yersinia* may hijack the vesicle secretion machinery by activating Cdc42 to enhance bacterial exocytosis.

To validate this assumption, we investigated whether CNF_Y_ influences Cdc42-GTP-mediated signaling pathways that are known to trigger Ca^2+^ mobilization and promote exocytosis. Exocytosis processes, particularly the late vesicle-plasma membrane fusion events, are triggered by the activation of adjacent Ca^2+^ channels - the IP_3_ receptors (IP_3_-R). IP_3_-R Ca^2+^ channels are opened upon inositol triphosphate (Ins(1,4,5)P_3_/IP_3_) binding, which is produced by phospholipase Cγ1 (PLC γ1) [35–37]. As constitutively active forms of Rac1 and Cdc42 have been shown to activate PLC γ1 and trigger vesicle exocytosis [36], we added inhibitors of PLCγ1 and the IP_3_-R in the Caco-2 monolayer system after infection with wildtype, Δ*cnfY* mutant +/- the empty vector (pV) or the complementation plasmid (p*cnfY*) to test their influence on bacterial egress. As shown in **Fig. 3D**, the number of egressed bacteria from the Caco-2 monolayer was strongly reduced upon the addition of the PLCγ1 or the IP_3_-R inhibitor for all the CNF_Y_-producing strains. Similar to the Cdc42 inhibitor, the number of cell-associated and intracellular CNF_Y_-expressing bacteria was not reduced upon this treatment; it was somewhat increased, likely due to the blockage of egress (Fig. **S6A-C**).

We also added specific activators of the two signaling molecules to support this finding. We tested whether they can enhance bacterial egress, particularly in infection with the low-egressing CNF_Y_-deficient mutant (Δ*cnfY*). In agreement with our previous results, activators for Cdc42/Rac-1, PLCγ1, and IP_3_-R were able to significantly increase the egress of the Δ*cnfY* mutant from the Caco-2 monolayer by about 4-6-fold (**Fig. 3E**) with comparable amounts of cell-associated bacteria (Fig. **S6D-F**). The addition of the activators did not significantly enhance the egress of the CNF_Y_-producing wild-type strain (**Fig. 3E**). This indicated that CNF_Y_-promoted activation of the signaling molecules was already at a high, near-maximal level.

Additionally, we quantified the IP_3_ concentration in Caco-2 monolayer cells at 3.5 hours post-infection, when maximal bacterial egress was observed. This was achieved by analyzing the accumulation of inositol monophosphate (IP_1_), a stable downstream metabolite of IP_3,_ upon the addition of LiCl, which blocks its metabolism to myo-inositol (**Fig. 3F**). IP_1_ is generated exclusively as a downstream product of IP_3_ following PLC γ1 activation (**Fig. 3F**) [38], and thus serves as a proxy for intracellular IP_3_ levels and PLC γ1 activation during bacterial egress. Using this assay, we found significantly lower levels of IP_1,_ and thus IP_3_, in the Caco-2 cells when the cell monolayer was infected with the Δ*cnfY* mutant compared to the CNF_Y_-positive wildtype and the complemented strain (**Fig. 3G**). Since none of these inhibitors affected the viability of Caco-2 cells (Fig. **S7**) or the growth of *Y. pseudotuberculosis* (Fig. **S8**), we concluded that the *Yersinia* toxin CNF_Y_ triggers the activation of a Cdc42-PLC1-IP3-IP3R cascade to enhance bacterial exit.

### *Y. pseudotuberculosis* egress involves SNARE-complex formation including VAMP-7 (v-SNARE), Stx4 (t-SNARE) and SNAP-23

The N-ethylmaleimide-sensitive factor (NSF) is crucial for vesicle and plasma membrane fusion, as well as exocytosis. They mediate the interaction of secretory vesicles with the plasma membrane attachment protein receptors found on the vesicles (initially named v-SNAREs) and the plasma membrane (initially named t-SNAREs) (**Fig. 4A**) [39]. These proteins form a stable complex via coiled-coil interactions (SNARE complex), pulling the membrane into proximity and loading the vesicles onto the plasma membrane [39,40]. This is followed by the priming process, rendering the vesicles competent for calcium-triggered fusion, a process shown to be promoted by the Rho GTPase Cdc42 [39,41]. Multiple subtypes of SNAREs were found to participate in (polarized) exocytotic processes in mammals (e.g. Syntaxins Stx-1, -2, -3, -4, the synaptosome-associated proteins SNAP-23, -25, the vesicle-associated proteins VAMP/-synaptobrevin-2, -7, -8, and also the late endosome/bacterial phagosome-associated proteins Syntaxin-7, -8, and VAMP-7, -8), depending on the implication of the subcellular compartments [39,40].

**Figure 4:**
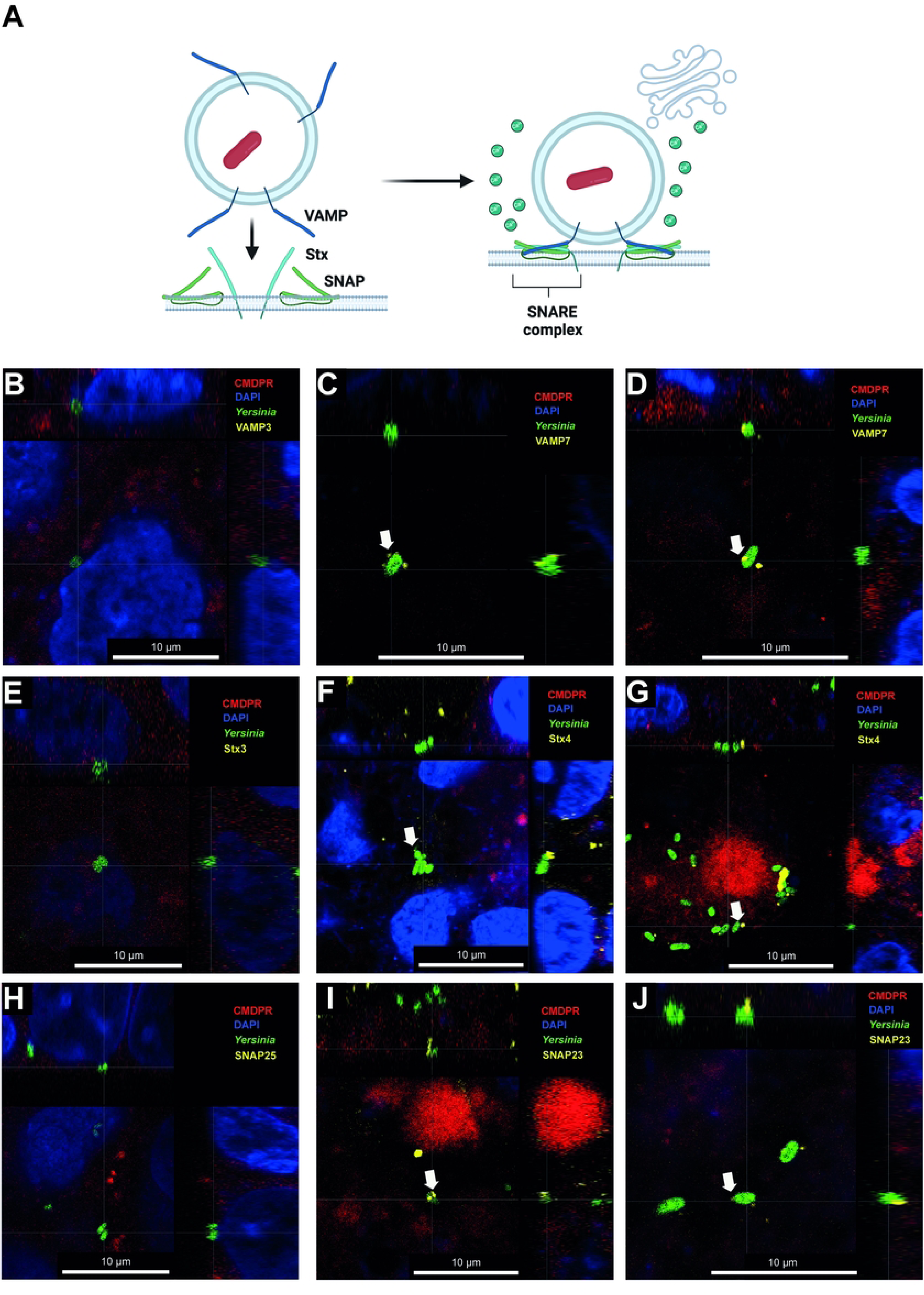
Association of the SNARE complex proteins VAMP3, VAMP7, Stx3, Stx4, SNAP23, and SNAP25 to intracellular *Yersiniae*. (**A**) Illustration of the VAMP-Stx-SNAP protein interaction for the formation of the SNARE complex initiating exocytotic processes. Created in BioRender. Dersch, **P.** (2026). (**B**-**J**) Caco-2 monolayer was infected with 2,5 x 10^6^ bacteria of *Y. pseudotuberculosis* wildtype YPIII pFU92 (*gfp*^+^). 3.5 h post-infection, the Caco-2 cell monolayers were analyzed by CLSM to detect *Yersinia* (GFP-green), the cell plasma membranes (CellMask Plasma Membrane Red-CMPR), the nuclei (DAPI-blue), and SNARE proteins (yellow) colocalized with the bacteria using antibodies against VAMP3 (**B**), VAMP7 (**C,D**), Stx3 (**E**), Stx4 (**F,G**), SNAP25 (**H**), and SNAP23 (**I,J**). Orthogonal XZ and YZ planes are shown.

To identify the molecular process involved in *Y. pseudotuberculosis* exit from host cells, we first examined the association of intracellular *Y. pseudotuberculosis* with VAMPs, which were most frequently found to participate in exocytotic processes using confocal laser scanning microscopy. GFP-expressing WT and Δ*cnfY* were used to infect the Caco-2 monolayer as described previously, and microscopic images were analyzed to colocalize *Yersinia*-containing vacuole (by the GFP) with VAMP3 and VAMP7. Early during transcytosis VAMP3-positive puncta structures were observed nearby intracellular *Y. pseudotuberculosis*, i.e., predominantly located beneath to the apical cell membrane of the Caco-2 cell monolayer, but not at bacteria at the basolateral membrane (**Fig. 4B**). In contrast, VAMP7 was identified as bound to the *Yersinia*-containing vacuoles close to the basolateral membrane at later time point of the infection (**Fig. 4C,D**). This indicates sequential recruitment and involvement of both v-SNARE molecules in intracellular trafficking, whereby VAMP7 appears to be more responsible for the docking and exit of *Yersinia*-containing vacuoles at the basolateral side.

We also tested whether and which members of the vesicle exocytosis machinery at plasma membranes (including Stx3 (**Fig. 4E**), Stx4 (**Fig. 4F-G**), SNAP25 (**Fig. 4H**), and SNAP23 (**Fig. 4I-J**) are recruited to the *Yersinia*-containing vacuole three hours post-infection, the time during transcytosis when the most frequent exit events occur. We found that Stx4 and SNAP23 were associated with the *Yersinia*-containing vacuole located at the basolateral membrane (**Fig. 4F-G, I-J**), whereas no association of Stx3 and SNAP25 was detected (**Fig. 4E, H**).

To further prove that both t-SNAREs, Stx4 and SNAP23, play a prominent role in the control of the *Yersinia*-containing vacuole fusion and content release after transcytosis, we constructed ΔStx4 and ΔSNAP23 knock-out derivatives of the Caco-2 cell lines, and an equivalent Caco-2 ΔStx3 cell line as a negative control (**Fig. 5A-C**). A ΔVAMP7 knock-out was not included in this analysis, as it has been shown that a deletion of the equivalent v-SNARE drastically affects *Yersinia* cell invasion and intracellular trafficking of the bacteria [42].

**Figure 5:**
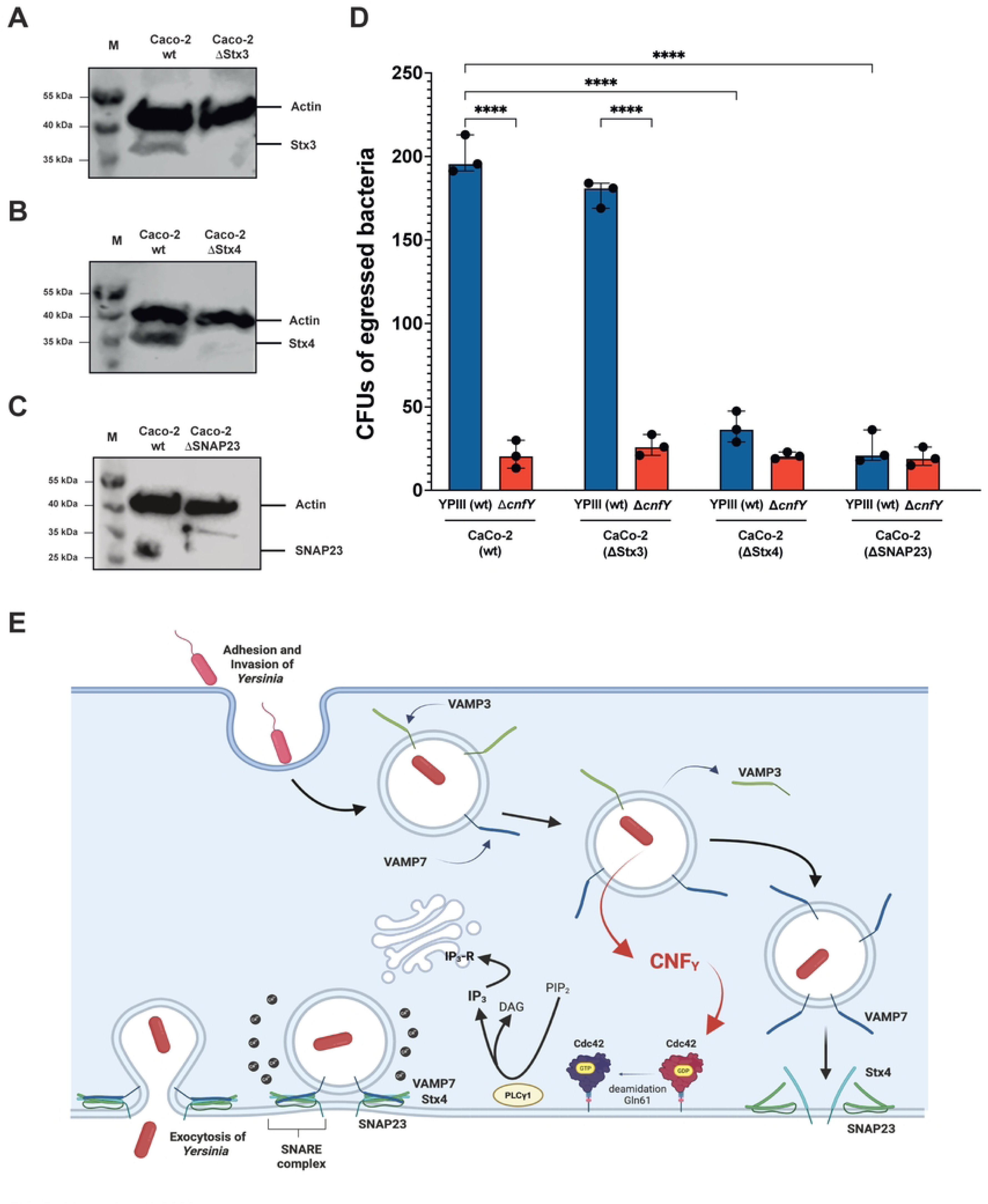
Role of SNARE complex proteins for *Yersinia* exocytosis and model of CNF_Y_-mediated increase of *Yersinia* cell egress. (**A-D**) Caco-2, and the isogenic cell lines Caco-2ΔStx3, Caco-2ΔStx4, Caco-2ΔSNAP23, and were seeded into transwells and cultivated until the TEER had reached > 400 Ω•cm². Whole-cell extracts were prepared and analyzed by Western blotting using antibodies specific to Stx3 (**A**), Stx4 (**B**), and SNAP23 (**C**), and actin as a loading control (**A-C**). M: protein marker. (**D**) Monolayers of wildtype Caco-2 cells (wt), and ΔStx3, ΔStx4, and ΔSNAP23 mutant derivatives were infected with *Y. pseudotuberculosis* wildtype strain YPIII (*cnfY*^+^) or YP147 (Δ*cnfY*). The CFUs of egressed bacteria were determined 3.5 h after infection. The mean +/- SEM of three independent biological replicates is shown. Significant differences were determined using a two-way ANOVA test with Tukey correction and indicated by asterisks (P-value: **** < 0.0001). (**E**) Illustration of the CNF_Y_-triggered activation of *Yersinia* egress through the induction of the Cdc42-dependent Ca^2+^-regulated exocytosis machinery. Created in BioRender. Dersch, **P.** (2026). Upon uptake of *Y. pseudotuberculosis,* the bacteria reside in a membrane-bound vacuole, to which the v-SNARE protein VAMP-3 is initially recruited. Later, during transcytosis, VAMP3 is replaced by VAMP7. The CNF_Y_ toxin is secreted by the intracellular bacteria (indicated by red arrows), leading to the activation of Cdc42 at the cell membrane by the deamidation of Gln61 of the Rho GTPase. This activates PLC-γ1, leading to the cleavage of phosphatidylinositol-4,5-bisphosphate (PIP_2_), producing inositol triphosphate (IP_3_) and diacylglycerol (DAG). IP_3_ interacts with the IP_3_ receptor (IP_3_-R) in the endoplasmic reticulum (ER), which triggers the activation of adjacent Ca^2+^ channels and the subsequent formation of a functional SNARE complex of VAMP7, Stx4, and SNAP25, and allows egress of *Yersinia* by exocytosis.

We first ensured that the deletions did not affect the development and integrity of the Caco-2 monolayer (Fig. **S9A-D**), did not cause cytotoxic effects (Fig. **S10A-C**), and had no negative influence on *Yersinia* cell adhesion and invasion (Fig. **S10D**). However, subsequent analyses of the exit rate revealed a significant reduction in the number of egressed bacteria from ΔStx4 and ΔSNAP23 Caco-2 cell monolayer compared to Caco-2 wildtype and the ΔStx3 knock-out cells, despite comparable *Yersinia* cell association rates (**Fig. 5D**, Fig. **S10D**). Taken together, this suggests that *Yersinia* exploits the regulated exocytotic machinery, specifically the VAMP7/Stx-4/SNAP-23 SNARE complex, to exit from intestinal epithelial cells and uses the CNF_Y_ toxin to stimulate this process by activating the Cdc42-PLC1-IP3-IP3R-Ca^2+^-pathway.

## Discussion

The ability to exit host cells is a fundamental step in the lifestyle of pathogenic bacteria, e.g., after intracellular proliferation or as the final step of transcytosis through epithelial cell barriers. Although bacterial cell escape is crucial for infecting neighboring cells and transmitting them to other tissues and susceptible hosts, it has received surprisingly little attention. To fill this knowledge gap, we studied the basolateral exit of *Y. pseudotuberculosis* from differentiated human gut epithelial cells following transcytosis using a standardized 2D monolayer model. We found that *Y. pseudotuberculosis* devises an escape strategy that harnesses specific host signaling pathways involved in vesicle release (exocytosis), leaving the host cell and, thus, the epithelial barrier intact. This is in contrast to many other intracellular bacteria, which disrupt host-cell membranes as they escape their host cells [2–6].

The *Yersinia*-containing vacuole (bacterial phagosome) is transported through the cell to the basolateral side of the epithelial cell monolayer, where it is docked and tethered to initiate its fusion with the plasma membrane, releasing the bacteria. After intestinal epithelial cell invasion, *Y. pseudotuberculosis* was found to reside in a single-membrane vacuole. In the early stages after cell entry, the *Yersinia*-containing vacuole (i.e., below the apical membrane) was strongly associated with the vesicle-associated membrane protein VAMP-3 and very little with VAMP-7. This finding aligns with a previous study, which showed that both v-SNAREs localize to intracellular *Yersiniae* residing in LC3-positive phagosomes in epithelial cells and macrophages [42]. During trafficking through the cell, a switch occurs, as all *Yersinia*-containing vacuoles, which are located close to the basolateral side of the cell, are exclusively associated with VAMP-7 (v-SNARE), and no VAMP-3 association is detectable. This suggests a different individual or sequential recruitment of both intracellular trafficking marker molecules, in agreement with studies reporting that VAMP3 is primarily present in early endosomes, whereas VAMP7 is predominantly found in late endosomes [43–46], which are known to be involved in exocytosis. Together, this suggests a potential role of the VAMPs in the control of YCV trafficking during transcytosis. However, their impact on intracellular bacterial trafficking during transcytosis remained unexplored.

A previous study analyzing the intracellular lifestyle of *Y. pseudotuberculosis* also identified both VAMPs on non-acidic *Yersinia*-containing vacuoles in HeLa cells. Here, VAMP3 favored the formation of single-versus double-membrane *Yersinia*-containing vacuoles, and VAMP7 was shown to be involved in the recruitment of LC3, implicated in autophagosome biogenesis [42,47]. Furthermore, VAMP3 and VAMP7 have been reported to be associated with vacuoles induced by other pathogens, including *Helicobacter pylori, Coxiella burnetti*, *Listeria monocytogenes,* and *Mycobacterium tuberculosis* [48–51]. As both v-SNARE molecules are key components of the vesicle fusion machinery, particularly of endosomes and lysosomes, they are assumed to play an important role in the biogenesis, maturation, and enlargement of bacterial-containing vacuoles, enabling persistence and replication within host cells. Only for *Listeria monocytogenes,* it was reported that VAMP3 is implicated in the bacteria-triggered remodeling of the host cell membrane into protrusions that internalize into neighboring cells and initiate cell-to-cell spreading [49].

After trafficking through the epithelial cells, the *Yersinia*-containing vacuole must dock and fuse with the basolateral membrane to release the pathogen into the extracellular space. This study shows that *Yersinia* hijacks the vesicle exocytosis machinery to initiate this process. Exocytosis of vesicles generally involves multiple coordinated steps, including several ‘priming’ events by a complex membrane fusion machinery and a fusion trigger, e.g., Ca^2+^, to proceed and complete the fusion [40].

Central to the final membrane fusion and exocytosis process is the formation of a core complex between the vesicular v-SNAREs (e.g., VAMP7) and the t-SNAREs (syntaxins and SNAP molecules) located at the target (basolateral) membrane, which is formed through coiled-coil structures [52]. Most of them are found in specific membrane compartments (e.g., Golgi, nucleus, endosomes, plasma membrane), promoting pairing and thus vesicle-fusion specificity [39,40]. Our colocalization studies and SNAP/Stx mutant analysis indicate that predominantly Stx4 and SNAP23 assemble with the VAMP7-decorated *Yersinia*-containing vacuole to form a functional SNARE complex to initiate the fusion of the vacuole and the plasma membrane, and the subsequent release of the bacteria (**Fig. 5E**). This combination is preferentially found in SNARE complexes, promoting late endosome fusion with the plasma membrane and vesicle exocytosis processes [53]. To initiate vesicle fusion, the VAMP protein on the vesicle interacts with the Stx and the SNAP proteins on the plasma membrane to form a four-helix bundle that zips up concomitant with the lipid bilayer fusion. This process creates curvature and causes lateral tension of the membranes, exposing the bilayer interior (hemifusion). This favors the contact between the distal membrane leaflets and subsequent formation of a fusion pore that expands and releases the vesicle content [39,40].

The SNARE proteins are usually kept inactive, but are quickly released to bind each other upon vesicle delivery and stimulation, e.g. by regulatory factors such as (i) tethering factors/complexes (e.g. octomeric exocyst/Rab effector complex (Sec3,5,6,8,10,15, Exo70,84), HOPS) and (ii) the synaptogamins, SM proteins (Vps33/subunit of the HOPS tethering complex) [40,52,54,55]. The tethering complexes mediate initial vesicle attachment to the plasma membrane prior to the function of the synaptogamins/SM proteins, which bind to different SNARE proteins to facilitate *trans* SNARE complex formation [40,52,54,55].

In recent years, it has also become clear that the Rho GTPase family plays a crucial role in regulating membrane trafficking and exocytosis [31,56,57]. The best-characterized members are RhoA, Cdc42, and Rac1, which can all be constitutively activated by the *Yersinia* toxin CNF_Y_ in murine macrophages [28]. However, the overall and local pools, as well as the activity, of the different Rho GTPases within cells vary significantly [57,58]. This may explain why the extent of CNF toxin-mediated activation of RhoA, Cdc42, and Rac1 is highly cell-type-dependent [28,59]. This study shows that *Y. pseudotuberculosis* CNF_Y_ predominantly activates the Rho GTPase Cdc42 in polarized human intestinal epithelial cells (Caco-2) and stimulates exocytosis by PLC γ1-dependent production of IP_3_, which leads to a rise of intracellular calcium known to initiate exocyst and SNAP complex formation (regulated exocytosis) [60] (**Fig. 5E**). This strongly indicates that *Y. pseudotuberculosis* uses the exocytosis machinery of vesicles to promote its cell exit during transcytosis of the intestinal barrier, and the CNF_Y_ toxin hijacks this process through activation of Cdc42. In fact, Cdc42 was shown to associate with the plasma membrane and was identified as the first Rho GTPase found to control intracellular secretory vesicle/endosome trafficking and exocytosis [57]. Activated Cdc42 was shown to interact directly with PLC1, thereby increasing intracellular IP3 levels. This initiates the release of Ca^2+^ from internal stores and an influx from the extracellular medium. The rise in intracellular Ca^2+^ is the primary trigger for exocytosis of secretory granules, e.g., in mast cells [36,61]. Several proteins associated with the vesicle fusion machinery were found to activate it upon Ca^2+^ binding, among them the synaptotagmins, which seem to play the most crucial role as Ca^2+^ sensors for exocytosis [62].

Besides activating the PLC1-IP_3_-Ca^2+^ pathway, it is possible that CNF_Y_-activated Cdc42 also enhances additional aspects of exocytosis. For instance, Cdc42 was shown (i) to interact with the exocyst subunits Sec3 and Exo70 in yeast, (ii) to bind to Syntaxin and VAMP2 in β-pancreatic cells to control insulin release and (iii) to manipulate N-WASP-Arp2/3-dependent actin polymerization and membrane tension and fusion pore dynamics, which was shown to be important for vesicle exocytosis by neuroendocrine cells [32,56,60,63,64]. Whether this holds for the bacterial egress process and how the bacteria trigger exocytosis without CNF_Y_ remains to be determined.

Our study indicates that, generally, only a few bacteria exit at the basolateral side of the intestinal epithelial layer. Although the CNF_Y_ toxin can significantly stimulate this process, the overall number of egressed bacteria is considerably lower than the initial inoculum that was internalized into the cells. The following events can explain this:

i. Multiple bacteria that invaded the intestinal epithelial layer at the apical side could be removed. In fact, a previous report using a human ileal enteroid-derived monolayer, including M cells, has shown that *Y. pseudotuberculosis*-mediated apical M cell extrusion occurs without inducing cell death or disrupting monolayer integrity [14]. The bacterial loss resulting from this event could not be quantified in our analysis because the antibiotic gentamicin was present on the apical side of our transwell model. Yet, we detected free-floating extruded cells with bacteria in our electron microscopy analysis during later stages of cell infection (Fig. **S2B**).
ii. Alternatively, the functional maturity of M cells in our experimental setup could be too low and may not reflect the cellular microenvironment found in the ileum overlying Peyer’s patches. However, M cells in our system expressed markers, such as GP2, indicative of maturity (**Fig. 1B,C**), and *Yersiniae* were bound to them in large numbers and were internalized (**Fig. 1E**). Similarly, many bacteria have been found to enter M cells of a human ileal enteroid-derived monolayer, which was differentiated in the presence of RANKL, TNF-α and CD137L to promote M cell development. However, although many *Yersiniae* invaded the M cells, only a tiny proportion (0.1-1%) was found in the basolateral chamber [14]. This suggests that epithelial cell barrier invasion is not the limiting step in transcytosis.
iii. Moreover, intracellular trafficking of the bacteria towards the basolateral side may allow host cell defense mechanisms to eliminate most bacteria residing in phagosomes. This assumption is supported by the frequent observation of ‘dying’ intracellular bacteria in our electron microscopy analysis (Fig. **S2A**). Notably, intracellular trafficking does not appear to be accelerated by the expression of the CNF_Y_ toxin, which would reduce the time bacteria spend encountering intracellular host-derived attacks, thereby enhancing their survival and cell egress. *Yersinia* exit was shown as a peak at 3.5 h, independent of the presence of the CNF_Y_ toxin. Interestingly, a linear peak has also been reported for the exit of *Porphyromonas gingivalis* at 6 hours post-infection from cultured immortalized human gingival epithelial cells [65], indicating that, once initiated, intracellular trafficking and exit dynamics are highly coordinated and timed/synchronized.

The fact that only a few bacteria seem to exit from the basolateral side of the intestinal epithelial layer indicates an extreme bottleneck for subsequent infection steps and would limit dissemination. In fact, a previous analysis using a mixture of green- and red-fluorescing *Yersiniae* revealed that the related gut pathogen *Yersinia enterocolitica* produces exclusively monoclonal microcolonies in the Peyer’s patches even at a high oral infection dose [66]. This indicated that single *Yersiniae* initiate abscess formation in the Peyer’s patches and deeper

tissues. Consistent with this observation, a more recent work using Sequence Tag-based Analysis of Microbial Populations (STAMP) with a barcoded population of *Y. pseudotuberculosis* showed that a minimal number of bacteria (between 10-20) initiate the colonization of the intestinal and systemic tissues, indicating a very tight bottleneck to colonization via the oral route [44]. Our results support this assumption, indicating that the limited egress of bacteria from the intestinal epithelial layer is the primary cause, rather than a few bacteria that survived the initial immune response in the lymphoid tissues.

As the expression of the CNF_Y_ toxin during transcytosis has been shown to significantly enhance the number of egressed bacteria, it is expected that the amounts of bacteria invading and colonizing the underlying Peyer’s patches will also increase. In a previous mouse infection study, in which we used a high dose of the wild-type and the *cnfY* mutant, we observed an equally high number of both strains in the Peyer’s patches after 24 h and the following days post-infection [28]. This would argue against a role of CNF_Y_ in this process. However, the initial high dose will likely allow some bacteria to exit the M cells and invade the underlying Peyer’s patches, even in the absence of CNF_Y_. Once the bacteria have invaded a Peyer’s patch, the host is shown to restrict subsequent infections [44,66]. The number of invaded bacteria and formed microcolonies remained constant, independent of the number of bacteria reaching the Peyer’s patches. We hypothesize that the exit-promoting role of CNF_Y_ may be critical in the very early infection phase, particularly when the initial infection dose is relatively low. However, testing this hypothesis with available mouse infection models is very challenging, as an early recovery of the bacteria from Peyer’s patches (e.g., a few hours post-infection), using a low infection dose (< 10^4^ bacteria), would be below the detection limit as only a few bacteria are expected to reach the underlying Peyer’s patches compared to the initial inoculum as outlined above.

Expression of CNF_Y_ during transcytosis could ensure that even a few bacteria that invaded M cells at low infection doses reach the underlying lymphoid tissues and initiate their systemic spread. On the contrary, it can also be an advantage to downregulate CNF_Y_ expression to reduce and/or delay the dissemination to other deeper tissues and organs, e.g., at higher infection doses, as the timing and efficiency of this process will determine whether the pathogen avoids or triggers the host’s innate and adaptive immune responses. In a previous study, we demonstrated that the absence of the CNF_Y_ toxin essentially prevents the typical *Yersinia*-triggered proinflammatory responses and associated tissue destruction (e.g., by enhancing interferon-mediated responses and tolerogenesis), facilitates the establishment of a commensal-like lifestyle, and drives *Y. pseudotuberculosis* into a persistent state [67]. The mechanism by which the reduction of bacteria exiting the M cells contributes to this process remains unclear and is currently under investigation. It is tempting to speculate that a low dose of invading CNF_Y_-negative bacteria may evade immediate recognition by residual innate immune cells, allowing the invaders more time to adapt to the new hostile tissue environment. However, additional processes are also likely to contribute to the establishment of persistence. For instance, the loss of CNF_Y_-mediated activation of Rho-GTPases in attacking immune cells, as well as the reduced injection of the Yop effectors by the bacteria in the absence of CNF_Y_, will have different effects on immune cell function and recruitment/migration compared to wild-type.

Altogether, this study reveals previously undescribed aspects of the transcytosis of enteric pathogens through the intestinal epithelial barrier, the step that allows the bacteria to escape from the intestine and spread to other tissues. Our findings define the molecular mechanism that triggers the egress of bacteria on the basolateral side of the intestinal cell layer. It reveals that the pathogen uses the Ca^2+^-regulated exocytosis machinery for its escape and highlights an unidentified role for Rho GTPase Cdc42-activating toxins in enhancing this process. Our results further suggest that transcytosis is a rare event that significantly limits bacterial dissemination, and illustrate how virulence factors, independent of the Type 3 secretion system, such as CNF_Y,_ contribute to pathogen fitness by optimizing early dissemination from the intestinal epithelium.

## Material and Methods

### Bacterial strains, media, and growth conditions

The *Y. pseudotuberculosis* wildtype strains IP32777, IP32954, IP2666, and YPIII, as well as their mutant derivatives, were used in this study (Table **S1**). To test the role of virulence factors for the *Yersinia* cell exit process, we used the pYV-cured derivative of YPIII (YP12), the YPIII Δ*cnfY* mutant (YP147), the IP2666 Δ*cnfY* mutants, and the complemented strains YPIII Δ*cnfY*/p*cnfY*^+^ (YP147 pJNS10) or the Δ*cnfY* mutant strain carrying the empty vector YPIII Δ*cnfY*/pV (YP147 pJNS11) [28]. For confocal microscopy, these strains were transformed with the GFP-expressing plasmid pFU92 [68].

*Yersinia* strains were grown in LB medium at 25°C or 37°C in LB (Luria Bertani) broth if not indicated otherwise. If necessary, antibiotics were added at the following concentrations: carbenicillin/ampicillin 100 µg/ml, chloramphenicol 30 µg/ml, and kanamycin 50 µg/ml. For Caco-2 monolayer infections, *Y. pseudotuberculosis* was grown at 25°C to the early stationary phase. The bacteria were pelleted and resuspended in DMEM Advanced (Gibco, #12491015) supplemented with 5% HyClone Fetal Bovine Serum (Cytiva, #1*6645735*), 200 mM L-glutamine (Gibco, #35050061), and 10 mg/ml human transferrin (Sigma-Aldrich, #T8158) and used to infect the Caco-2 cells at an MOI of 50.

### Cell culture, intestinal cell monolayer system, and growth conditions

Caco-2 cells were cultured in DMEM Advanced (Gibco, #12491015) supplemented with 5% HyClone Fetal Bovine Serum (Cytiva, #16645735), 200 mM L-glutamine (Gibco, #35050061), and 10 mg/ml human transferrin (Sigma-Aldrich, #T8158) and grown in a cell incubator at 37°C, 5% CO_2_. For egress assays and electron microscopy, 3 µm pore-size translucent PC-inserts (Corning, #3415) were used. For confocal microscopy, 0.4 µm pore-size transparent PET inserts (Corning, #3470) were used. Inserts were coated with extracellular matrix (Tebu, Offenbach, #MTSIN201) diluted 1:100 in PBS in a 24-well plate. The growth media of the upper and basal chambers were refreshed every two days after seeding. To ensure the integrity of the monolayer throughout the experiment, transepithelial electrical resistance (TEER) was continuously measured using a CellZScope (NanoAnalytics). The Caco-2 cells were grown until they reached a TEER > 400 Ω·cm^2^ (approximately 10 days).

After the experiment, permeability assays were performed using 40 kDa TRITC-Dextran (Invitrogen, #D1842). The transwells were incubated with 200 µg/ml TRITC-dextran for 30 min, and the permeability of the cells was analyzed by detecting the TRITC-mediated fluorescence in the basolateral chamber. As a positive control for the permeability assay, a monolayer in an uninfected transwell was destroyed using 0.1% Triton X-100. All transwells with increased TRITC signal, compared to uninfected transwells or reduced TEER (TEER < 300 Ω·cm^2^) on the day or after infection, were excluded from the transcytosis assay.

### Construction of strains

To construct IP2666 Δ*cnfY*, plasmid pJNS5 was transformed into S17λpir and transferred into *Y. pseudotuberculosis* IP2666 via conjugation. Chromosomal integration of the plasmid was selected by plating on LB supplemented with kanamycin. Single colonies were tested for correct plasmid integration via colony PCR using primer pair III711/III713. Mutants were subsequently grown on LB agar plates containing 10% sucrose. Single colonies were tested for growth on carbenicillin and sequenced using primer pair III711/III713.

### Infection of the Caco-2 monolayer and transcytosis assay

Caco-2 cells grown in transwells were infected apically at an MOI of 50 and incubated for up to 10 hours. After 2-4 hours post-infection, the media in the upper chamber were removed and replaced with a medium containing gentamicin (to kill extracellular bacteria on the apical side of the Caco-2 monolayer), with or without inhibitors/activators of cell signaling pathways/molecules with the following concentration of the inhibitors for RhoA (30 µM, Rhosin, Sigma-Aldrich, #555460), Rac-1/Cdc42 (2.5 µM ML141, MedChemExpress, #HY-12755), PLCγ-1 (5 µM, U-73122, MedChemExpress, #HY-13419), IP_3_-R (40 µM 2-APB, MedChemExpress, #HY-W009724), and the activators of Cdc42/Rac1 (1 unit/ml, Rac1/Cdc42 activator II, Cytoskeleton, Inc., #CN02-A), PLC γ-1 (15 µM m-3M3FBS, MedChemExpress, #HY-19619), and IP_3_-R (30 µM D-myo-Inositol-1,2,4,5-tetraphosphate, sodium salt, Santa Cruz Biotechnology, #sc-362076). The medium of the basolateral chamber was collected at the indicated time points post-infection, replaced with fresh medium, and the removed medium was plated on LB agar plates to determine the CFUs of bacteria that had egressed from the Caco-2 cell monolayer.

### Adhesion and invasion assay

Adhesion/cell association, as well as bacterial penetration into the Caco-2 cell monolayer, were assessed using adherence and gentamicin protection assays, as described [23,69]. Bacteria were added to the cell monolayer (5 x 10^4^ Caco-2 cells in transwells) at an MOI of 50. For the adherence/cell association assay, the samples were incubated for 3 hours before cell lysis with 0.1% Triton X-100. To determine the number of intracellular bacteria at this time point, gentamicin was added and incubated for at least 60 minutes to kill extracellular bacteria. The surviving intracellular bacteria were released with 0.1% Triton X-100, and the bacteria in the Triton X-100-treated samples were titrated on LB plates.

### Cell viability assay

Lactate dehydrogenase (LDH) release from damaged cells, as determined with an LDH activity assay kit (Promega, #G1780), was used to assess Caco-2 cell viability. Caco-2 cells were seeded in transwells and grown as described above. Before the assay, the cells of the monolayer were washed with PBS and incubated in fresh cell culture medium. Subsequently, the LDH detection solution of the assay, containing NAD and a NADH-specific probe, was added to the cell medium in the apical chamber. Released LDH was then detected as described by the manufacturer after 1 hour of incubation. Briefly, released LDH in the supernatant reduces NAD in the detection solution, followed by the interaction with a probe, which was detected by the absorbance at 490 nm of the cell supernatant using a plate reader (CLARIOstar Plus).

### Construction of mutant Caco2 cell derivatives

For the construction of Caco-2 deletion cell lines, CRISPR/Cas9 KO and HDR plasmids from Santa Cruz Biotechnology were used. For Caco-2 ΔStx3, the Syntaxin 3 CRISPR/Cas9 KO Plasmid (h) (Santa Cruz Biotechnology, #sc-403466) and Syntaxin 3 HDR Plasmid (h) (Santa Cruz Biotechnology, sc-403466-HDR) were used. For Caco-2 ΔStx4, the Syntaxin 4 CRISPR/Cas9 KO Plasmid (h) (Santa Cruz Biotechnology, #sc-402879) and Syntaxin 4 HDR Plasmid (h) (Santa Cruz Biotechnology, #sc-402879-HDR) were used. For Caco-2 ΔSNAP23, SNAP23 CRISPR/Cas9 KO Plasmid (h) (Santa Cruz Biotechnology, #sc-417781) and the SNAP23 HDR Plasmid (h) (Santa Cruz Biotechnology, #sc-417781-HDR) were used.

2 x 10^5^ Caco-2 cells were seeded into a 6-well tissue culture plate. One day after seeding, transfection with the plasmids was performed as described by the manufacturer. Briefly, 1.5 µg of KO and HDR Plasmid DNA was diluted in 150 µl of Plasmid Transfection Medium (Santa Cruz Biotechnology, #sc-108062), and 10 µl of UltraCruz Transfection Reagent (Santa Cruz Biotechnology, #sc-395739) was added to 140 µl of Plasmid Transfection medium (Santa Cruz Biotechnology, #sc-108062), and incubated for five minutes at room temperature. The KO and HDR plasmid-containing solution was added dropwise to the solution containing UltraCruz Transfection Reagent, and the mixture was incubated at room temperature for 30 minutes. The resulting mixture was added dropwise to the wells and incubated for 48 hours post-transfection. Subsequently, the growth media were changed to media containing 2.5 µg/ml of puromycin (Santa Cruz Biotechnology, #sc-108071) and incubated for 5 days with media exchange every 2 days. Gene disruption of the knock-out cell lines was confirmed by Western blotting of whole-cell extracts using the primary polyclonal antibodies against Stx3 (1:1000, Invitrogen, #PA5-51691), Stx4 (1:000, Invitrogen, #PA5-51560), SNAP23 (1:1000, Invitrogen, #PA1-738) and SNAP25 (1:1000, Invitrogen, #PA1-740), and the secondary Goat Anti-Rabbit HRP antibody (1:5000, Jackson, #111035003) for detection. For loading control, the β-Actin mouse mAb was used (1:1000, Sigma, #A5441).

### Small GTPase Activation Assay

Activation of different small GTPases (RhoA, Rac1, Cdc42) in Caco-2 cells was tested with the RhoA/Rac1/Cdc42 Activation Assay Combo Kit (Cell Biolabs Inc., #STA-405) as described by the manufacturer. Briefly, Caco-2 cells were grown for 10 days on transwell plates and infected with the indicated *Y. pseudotuberculosis* strains with an MOI of 50. After 3.5 hours, cells were harvested, lysed, and cell extracts were prepared. Rhothekin RBD beads for RhoA and PAK PBD Agarose beads for Cdc42 and Rac1 pull-down were added to the cell extract and incubated for 1 hour at 4°C and 300 rpm. The activated Rho GTPases and the total amount of GTPases were identified and visualized by SDS-PAGE according to the manufacturer’s instructions. The beads were pelleted by centrifugation, washed with Assay Buffer (Cell Biolabs Inc., #STA-405), and resuspended in SDS-PAGE sample buffer. The proteins in the samples were separated by SDS-PAGE. The activated Rho GTPases in the samples were detected by Western blotting using the Anti-RhoA, Anti-Cdc42, and Anti-Rac1 antibodies (1:1000, Cell Biolabs Inc., #STA-405), and the goat anti-mouse-HRP secondary antibody (1:5000, Jackson, #115-035-044).

### IP_3_ Quantification

The quantification of accumulated D-myo-inositol 1-phosphate (IP_1_) was used as a proxy for the IP_3_ concentration, as IP_1_ is produced solely from IP_3_ by PLC1. It accumulates as a stable downstream metabolite of IP_3_ upon adding LiCl, which blocks its metabolism to myoinositol. For quantification, we used the IP-One ELISA Kit (Revvity, #72IP1PEA) according to the manufacturer’s protocol.

Caco-2 cells were seeded in a 24-transwell plate at 4 x 10^5^ cells/well and infected with the indicated *Y. pseudotuberculosis* strains at an MOI of 50. 50 mM LiCl was added to the cells at the time of infection with the bacteria to block the degradation of IP_1_. After 3.5 hours, cells were washed with PBS and lysed for 30 minutes with 50 µl of 2.5% lysis reagent. 50 µl of supernatant was transferred into an ELISA-coated plate. 25 µl of IP_1_-HRP conjugate and 25 µl of Anti-IP_1_ Mab (Revvity, #72IP1PEA) were added, and the plate was incubated for 3 hours at room temperature at 300 rpm. The supernatants were removed, and the plate was washed several times with the wash solution (Revvity, #72IP1PEA). Afterwards, it was incubated at room temperature in the dark with 100 µl TMB (3,3’,5,5’-tetramethylbenzidine) substrate solution, a chromogen to detect horseradish peroxidase (HRP) enzyme activity. After 30 minutes, the reaction is stopped with 100 µl of stop solution, and the absorbance is measured at 450 nm with correction at 620 nm.

### Preparation of cell extracts, gel electrophoresis, and western blotting

For immunological detection of specific proteins of Caco-2 cells, cell extracts of (un)treated cells were prepared and separated on 12-15% SDS polyacrylamide gels [70]. The proteins from SDS-PAGE were transferred onto an Immobilon PVDF membrane (Millipore) by electroblotting. The membranes were then blocked with 3% non-fat dry milk in TBS-T (Tris-buffered saline with 0.1% Tween 20) for 1 hour, and probed with the respective primary and secondary antibodies, diluted in 3% non-fat dry milk in TBS-T, as described [71]. For visualization, the membranes were developed by using the Western Lightning ECL II Kit (Cell Signalling, #6883). The proteins of interest were detected using LI-COR imaging system (Biosciences).

### Immunofluorescence staining and confocal laser scanning microscopy (C-LSM)

After treatment, Caco-2 cells in the transwell were washed with PBS and fixed in 4% formaldehyde for 10 minutes at room temperature. Subsequently, cells were permeabilized with 0.1% Triton in PBS for five minutes. Cells were then blocked in 3% BSA in PBS for one hour at room temperature and incubated overnight at 4°C with: (1) DAPI (1:1000, Invitrogen, #62248) to stain the nuclei, (2) CellMask Plasma Membrane Red (CMPR) (1:1000, Invitrogen, #C10046) to stain the cell membranes, (3) primary antibody against Glycoprotein-2/Anti-GP2 (1:100, Invitrogen, #PA5-42593) to test the development of M cell in the monolayer, (4) primary antibodies against VAMP3 or VAMP7 (1:100, Invitrogen, #MA5-34747 and # PA5-116892) to test colocalization of the VAMPs with the *Yersinia*-containing vacuole membrane, and (5) primary antibodies against Stx3, Stx4, SNAP23, or SNAP25 (1:100, Invitrogen, #PA5-51691, #PA5-51560, #PA1-738, and #PA1-740) to test colocalization of the t-SNARES.

After that, the cells were washed with PBS and incubated with secondary antibody, Goat anti-mouse IgG Alexa Fluor 568 (Invitrogen, #A-11011), at room temperature for 2 hours, followed by PBS washing. The transwell membranes were cut from inserts and mounted on slides using CitiFluor AF1 mounting media (EMS, #E17970-25). Coverslips were sealed with nail polish.

Samples were visualized using a laser scanning confocal microscope (Stellaris 5 WLL, Leica). Technical details on imaging are provided in Supplementary Note **A**. The Confocal microscopy images were adjusted and exported using LASX software (Version 4.7.0.28176). Technical information about adjustment, analysis, and exportation is provided in Supplementary Note **B**.

### Electron microscope

The 2D Caco-2 monolayers in transwells were fixed with 2% glutaraldehyde (Polysciences, #01909) in PBS (pH 7.2) for 10 minutes and then incubated for 2 hours at room temperature. The samples were stored at 4°C until further processing. After fixation, the samples were washed three times with PBS (pH 7.2), post-fixed in 1% osmium tetroxide (Electron Microscopy Sciences, #19100) in PBS at room temperature for one hour, and washed with PBS. Samples were gradually dehydrated in increasing ethanol concentrations (50%, 70%, 90%, 96%, 99.8%) for 20 minutes at each step at RT. Block contrasting was performed with saturated uranyl acetate (Polysciences, #21447) in 70% ethanol during gradual dehydration. After incubation with 100% propylene oxide (Serva, #33715) twice for 5 minutes, subsequent incubation embedded the samples with Epoxy resin components 2,4,6-Tris(dimethylaminomethyl)phenol, 2-Dodecenylbernsteinsäureanhydrid, Glycidether 100, and Methylnadinsäureanhydrid (Serva, #36975, #20755, #21045, and #29452) mixed with propylene oxide (1:1, 2:1 each step 2 hours RT), epoxy resin 100% (2x 2 hours RT), and polymerized in fresh epoxy resin at 60°C for 72 hours. Ultrathin sections (60 nm) were generated using a Leica EM UC7 ultramicrotome. Sections were poststained with 4% uranyl acetate (Polysciences, #21447) in 25% ethanol and ‘Reynold’s lead citrate’ with the components Tri-sodium citrate dihydrate and Lead (II) citrate (Merck, #106448 and Polysciences #00785). Samples were analyzed at 80 kV using a FEI Tecnai 12 electron microscope (FEI, Olympus). Images of selected areas were documented and analyzed using a Veleta 4 K CCD camera (Emsis) and Tecnai software (FEI/Thermo Fisher Scientific). Scale bars are indicated in the respective images.

### Quantifications and statistics

Statistical analyses were performed using GraphPad Prism Software version 9.4.1. Statistical tests used are described in individual figure legends, where * p < 0.05, ** p < 0.01, *** p < 0.001, **** p < 0.0001, and ns = no significance for all figures and supplemental tables.

## Acknowledgments

We thank the master’s students Dana Unländer and Luka Ressmann for experimental support, and Britta Körner for cell culture assistance.

## Resource availability

Lead contact

For further information or access to the pathogens, reagents, or mice used in the study, please contact the lead contact, Petra Dersch.

## Materials availability

Bacteria and cell lines generated in this study, as well as the vectors used to create the bacterial mutants, are available upon request.

## Data and code availability

All raw data, including any additional information required to reanalyze the data reported in this paper, are available on request. This manuscript does not report original code. Upon request, the lead contact can also provide any additional information required to reanalyze the (imaging) data reported in this paper.

## Author contributions

**Conceptualization and outline of the study:** Petra Dersch, Christian Rüter

**Data curation:** Christopher Margraf

**Formal analysis:** Christopher Margraf

**Funding acquisition:** Petra Dersch

**Investigation:** Christopher Margraf, Joyleen Fernandes, Lilo Greune, Sandra Heissler, Johanna Sibbel

**Experimental design:** Christopher Margraf, Petra Dersch, Christian Rüter

**Methodology:** Christopher Margraf, Paweena Wessel, Samriti Sharma

**Project administration:** Petra Dersch

**Software:** not applicable

**Resources:** Petra Dersch

**Supervision:** Petra Dersch, Christian Rüter

**Validation:** Christopher Margraf, Petra Dersch

**Visualization:** Christopher Margraf, Lilo Greune

**Writing – original draft:** Christopher Margraf and Petra Dersch drafted the original version

**Writing – review & editing:** Petra Dersch, Christopher Margraf, Christian Rüter, Paweena Wessel, Joyleen Fernandes, Samriti Sharma, with input from all listed authors

## Supplementary Figures

**Figure S1: Analysis of the integrity of the Caco-2 monolayer established for the analysis of bacterial transcytosis.** Approximately 10^4^ Caco-2 cells were seeded into transwell inserts (3 µm pore size) and cultivated until the TEER had reached > 300 Ω•cm². (**A**) 40 kDa TRITC-dextran permeability assays were used to assess the monolayer integrity. TRITC-dextran was added to the apical chamber. At the indicated time points, the medium of the basolateral chamber was removed, and TRITC fluorescence was determined. For positive control, the Caco-2 cell monolayer was treated with 0.1% Triton X-100. The mean +/- SEM of three independent biological replicates is shown. Significant differences were determined using a Kruskal-Wallis or ordinary one-way ANOVA test with Tukey correction and indicated by asterisks (P-value: ** < 0.01, **** < 0.0001). (**B**) The transepithelial electrical resistance (TEER) of the Caco-2 cells was measured continuously after seeding the cells into the transwell plates and after infection with *Y. pseudotuberculosis* wildtype strain YPIII (wt), and the Δ*cnfY* mutant strain YP147 (YPIII Δ*cnfY*) +/- the *cnfY*^+^ complementation plasmid pJNS10 (p*cnfY*^+^). The change of media and the start of the infection are indicated. (**C**) Analysis of the cell viability of the Caco-2 cells in the monolayer over the course of the infection. The Caco-2 cell monolayer was infected with an MOI of 50 with *Y. pseudotuberculosis* wildtype strain YPIII (wt), and the virulence plasmid-cured strain YP12 (ΔpYV). After the indicated time points, the viability of the Caco-2 cells in the monolayer was analyzed by the LDH release assay. For positive control, the Caco-2 cell monolayer was treated with 0.1% Triton X-100. The relative LDH release was determined as the percentage of LDH release in infected samples relative to the positive control, set at 100%. The mean +/- SEM of three independent biological replicates is shown.

**Figure S2: Electron microscopy of an intracellular dying *Yersinia* cell and exfoliating cells of the Caco-2 monolayer.** Caco-2 cells were infected with the *Y. pseudotuberculosis* wildtype strain YPIII. 3.5 hours post-infection, the cells were harvested and prepared for transmission electron microscopy. Images of (**A**) an intracellular dying *Yersinia* cell in a membrane-enclosed vacuole, and (**B**) exfoliating cells infected with *Yersiniae* are shown.

**Figure S3: Comparative analysis of *Yersinia*-mediated cell toxicity, cell adhesion, and invasion of strains with and without expression of CNF_Y_.**

Caco-2 cell monolayers in transwells were infected with 2.5 x 10^6^ bacteria of the *Y. pseudotuberculosis* wildtype strain YPIII (*cnfY*^+^), and the Δ*cnfY* mutant strains YP147 (YPIII Δ*cnfY*) +/- the empty vector pJNS11 (pV) and the *cnfY*^+^ complementation plasmid pJNS10 (p*cnfY*). (**A**) 3 hours (h) post-infection (pi), the Caco-2 cells were washed and lysed to determine the number of cell-associated bacteria. For the analysis of intracellular bacteria, gentamicin was added 3 hours post-infection. After an additional 2 hours of incubation, the Caco-2 cells were washed and lysed. The number of cell-associated and intracellular bacteria in the cell extracts was then determined by plating onto LB agar. The mean +/- SEM of three independent biological replicates is shown. Significant differences were determined using a two-way ANOVA test with Tukey correction and indicated by asterisks (P-value: ** < 0.01, *** < 0.001, **** < 0.0001). (**B**) In a parallel set-up, 3 hours post-infection, the infected Caco-2 cell monolayer was analyzed by the LDH release assay (3 h pi), or the Caco-2 cell monolayer was washed and incubated in the presence of gentamicin for two additional hours before the LDH release assay was performed. For positive control, the Caco-2 cell monolayer was lysed with 0.1% Triton X-100. The relative LDH release was determined as the percentage of the LDH release of infected samples relative to the positive control, set at 100%. The mean +/- SEM of three independent biological replicates is shown.

**Figure S4: Comparative analysis of cell adhesion, invasion, and triggered cell toxicity by *Y. pseudotuberculosis* with and without expression of CNF_Y_.**

Caco-2 cell monolayers were infected with different *Y. pseudotuberculosis* isolates (IP32777, IP32953, and IP2666, of which IP2666 encodes a functional *cnfY* gene) and the isogenic Δ*cnfY* mutant derivative IP2666 Δ*cnfY* wildtype. (**A**) 3 h post-infection, the Caco-2 cells were washed and lysed to determine the number of cell-associated bacteria. For the analysis of intracellular bacteria, gentamicin was added 3 hours post-infection. After an additional 2 hours of incubation, the Caco-2 cells were washed and lysed. The number of cell-associated and intracellular bacteria in the cell extracts was then determined by plating onto LB agar. The mean +/- SEM of three independent biological replicates is shown. The significant differences were determined using a two-way ANOVA test with Tukey correction and indicated by asterisks (P-value: * < 0.05, ** < 0.01). (**B**) In a parallel setup, 3 hours post-infection, the infected Caco-2 cell monolayer was analyzed using the LDH release assay (3 h pi). Alternatively, the Caco-2 cell monolayer was washed and incubated with gentamicin for 2 additional hours before the LDH release assay was performed. For positive control, the Caco-2 cell monolayer was lysed with 0.1% Triton X-100 The relative LDH release was determined as the percentage of the LDH release of infected samples relative to the positive control, set at 100%. The mean +/- SEM of three independent biological replicates is shown.

**Figure S5: Analysis of the influence of Rho GTPase RhoA on bacterial egress.**

Caco-2 monolayer was infected with 2.5 x 10^6^ bacteria of the *Y. pseudotuberculosis* wildtype strain YPIII (*cnfY*^+^), YP147 (YPIII Δ*cnfY*) +/- the empty vector pJNS11 (pV), or the *cnfY*^+^ complementation plasmid pJNS10 (p*cnfY*). (**A**) 3 hours post-infection, gentamicin was added to kill extracellular bacteria, and an inhibitor of the Rho GTPase RhoA (Rhosin) was added. After an additional 30 minutes of incubation, the medium from the basolateral chamber was collected and plated on LB plates to determine the CFUs of egressed bacteria. The mean +/- SEM of three independent biological replicates is shown. A two-way ANOVA test with Tukey correction showed no significant differences. (**B**,**C**) Caco-2 monolayer was infected with *Y. pseudotuberculosis* wildtype YPIII (*cnfY*^+^), the *Yersinia* virulence-negative strain (ΔpYV), the *cnfY*-negative mutant strain YP147 (Δ*cnfY*) without or with *cnfY*^+^ complementation plasmid pJNS10 (p*cnfY*^+^). After 3.5 h post-infection, infected Caco-2 cells were lysed, cell extracts were prepared, and activation of the different Rho GTPases was tested by the isolation of the GTP-bound form by pull-downs with Rho-GTPase-binding agarose beads and Western blotting using Rho GTPase-specific antibodies directed against RhoA (**B**) or Rac1 (**C**). M: Protein size marker. As negative and positive controls, high concentrations of GDP and GTP (GTPγS) were added to uninfected whole-cell extract samples, and purified RhoA (**B**) or Rac1 (**C**) proteins (+) supplied by the detection assay were loaded onto the SDS gel to ensure antibody binding and recognition. Equal concentrations of extracts were used for pull-down assays (**B, C**), which were assessed by Western blotting with Actin antibodies.

**Figure S6: Influence of GTPase, PLC γ-1, and IP3-R inhibitors and activators on Caco-2 cell monolayer adhesion and invasion.**

Caco-2 monolayer was infected with 2.5 x 10^6^ bacteria of the *Y. pseudotuberculosis* wildtype strain YPIII (wt), YP147 (Δ*cnfY*) +/- the empty vector pJNS11 (pV), or the *cnfY*^+^ complementation plasmid pJNS10 (p*cnfY*). 3 hours post-infection, inhibitors (Cdc42/Rac1 (ML141) (**A**), phospholipase C (PLC γ-1) (U-73122) (**B**), and Ca^2+^ channel IP3-R (2-APB) (**C**)), or activators (Cdc42/Rac1 (Activator II) (**D**), PLC γ-1 (m-3M3FBS) (**E**), and IP3-R (Myo-Inositol) (**F**)) were added to the apical side of the monolayer. After an additional 30 min, Caco-2 cells were washed and lysed to determine the number of cell-associated bacteria. For the analysis of invaded bacteria, gentamicin was added three hours post-infection, together with the inhibitors and activators. After an additional two hours of incubation (5 hours post-infection), the Caco-2 cells were washed and lysed. The number of cell-associated and intracellular bacteria in the cell extracts was then determined by plating onto LB agar. The mean +/- SEM of three independent biological replicates is shown. A two-way ANOVA test with Tukey correction was used for statistical analysis. P-value: * < 0.05.

**Figure S7: Analysis of cell viability in the presence of GTPase, PLC γ-1, and IP3-R inhibitors and activators.** Inhibitors (**A**) or activators (**B**) of the Rho GTPase Cdc42, PLC γ-1, and IP3-R were added to the Caco-2 cell monolayer and incubated for 3 and 5 hours, respectively, covering the time points in which bacterial cell-associated and egress were tested. After the indicated time points, the media were withdrawn and analyzed using an LDH release assay. For positive control, the Caco-2 cell monolayer was lysed with 0.1% Triton X-100. The relative LDH release was determined as the percentage of the LDH release in individual samples relative to the positive control, which was set at 100%. The mean +/- SEM of three independent biological replicates is shown.

**Figure S8: Influence of GTPase, PLC γ-1 and IP3-R inhibitors and activators on the viability and growth of the Caco-2 cells or *Y. pseudotuberculosis***

*Y. pseudotuberculosis* wildtype YPIII (*cnfY*^+^), the YP147 (YPIII Δ*cnfY*) +/- the empty vector pJNS11 (pV) and the *cnfY*^+^ complementation plasmid pJNS10 (p*cnfY*^+^) were diluted 1:1000 from overnight cultures and grown in LB without (**A**) or in the presence of the different, indicated Rho GTPase (**B,C**), PLC γ-1 (**D**) and IP3-R inhibitors (**E**) and activators (**F,G,H**) at 37°C. Bacterial growth was monitored by measuring optical density at 600 nm (OD_600)_ over 6 hours. The mean +/- SEM of three independent experiments is shown.

**Figure S9: Analysis of the Caco-2 ΔStx3, ΔStx4, and SNAP23 knock-out cells.**

(**A**) Caco-2 cells (wt) and the the Caco-2ΔStx3 (**B**), Caco-2ΔStx4 (**C**) and Caco-2ΔSNAP23 (**D**) mutants were seeded into transwells (3 µm pore size) and cultivated for approximately 10 days (5 x 10^4^ cells per well), before infection with 2.5 x 10^6^ bacteria of the *Y. pseudotuberculosis* wildtype YPIII (*cnfY*^+^) strain. The transepithelial electrical resistance (TEER) of the different Caco-2 cell variants was measured continuously after seeding the cells into transwell plates and culturing until the TEER reached > 300 Ω•cm². The change of media and the start of the infection are indicated.

**Figure S10:** Analysis of the cell adhesion, invasion, and triggered cell toxicity by *Y. pseudotuberculosis* with and without CNF_Y_ expression with Caco-2 ΔStx3, ΔStx4, and SNAP23 knock-out cells. The monolayer of Caco-2 cell wildtype and mutants was infected with 2.5 x 10^6^ bacteria of the *Y. pseudotuberculosis* wildtype strain YPIII (wt) (MOI of 50). 3 h post-infection (pi), the media of the monolayer were withdrawn for analysis and replaced by media containing gentamicin. After an additional 2 h, the media were again withdrawn. Subsequently, all media were analyzed using an LDH release assay. For positive control, the Caco-2 cell monolayer was lysed with 0.1% Triton X-100. The relative LDH release was determined as the percentage of the LDH release in the individual samples with (**A**) Caco-2 ΔStx3, (**B**) ΔStx4, and (**C**) SNAP23 cells relative to the positive control, which was set at 100%. (**D**) In a parallel set-up, Caco-2 cells were washed and lysed 3 hours post-infection to determine the number of cell-associated bacteria. For the analysis of invaded bacteria, gentamicin was added 3 hours post-infection (3 h pi), and after an additional 2 hours incubation (5 h pi), the Caco-2 cells were washed and lysed. The number of cell-associated and intracellular bacteria in the cell extracts was then determined by plating onto LB agar. The mean +/- SEM of three independent biological replicates is shown. Significant differences were determined using a two-way ANOVA test with Tukey correction and indicated by asterisks (P- value: * < 0.05, ** < 0.01, *** < 0.001).

**Table S1:**
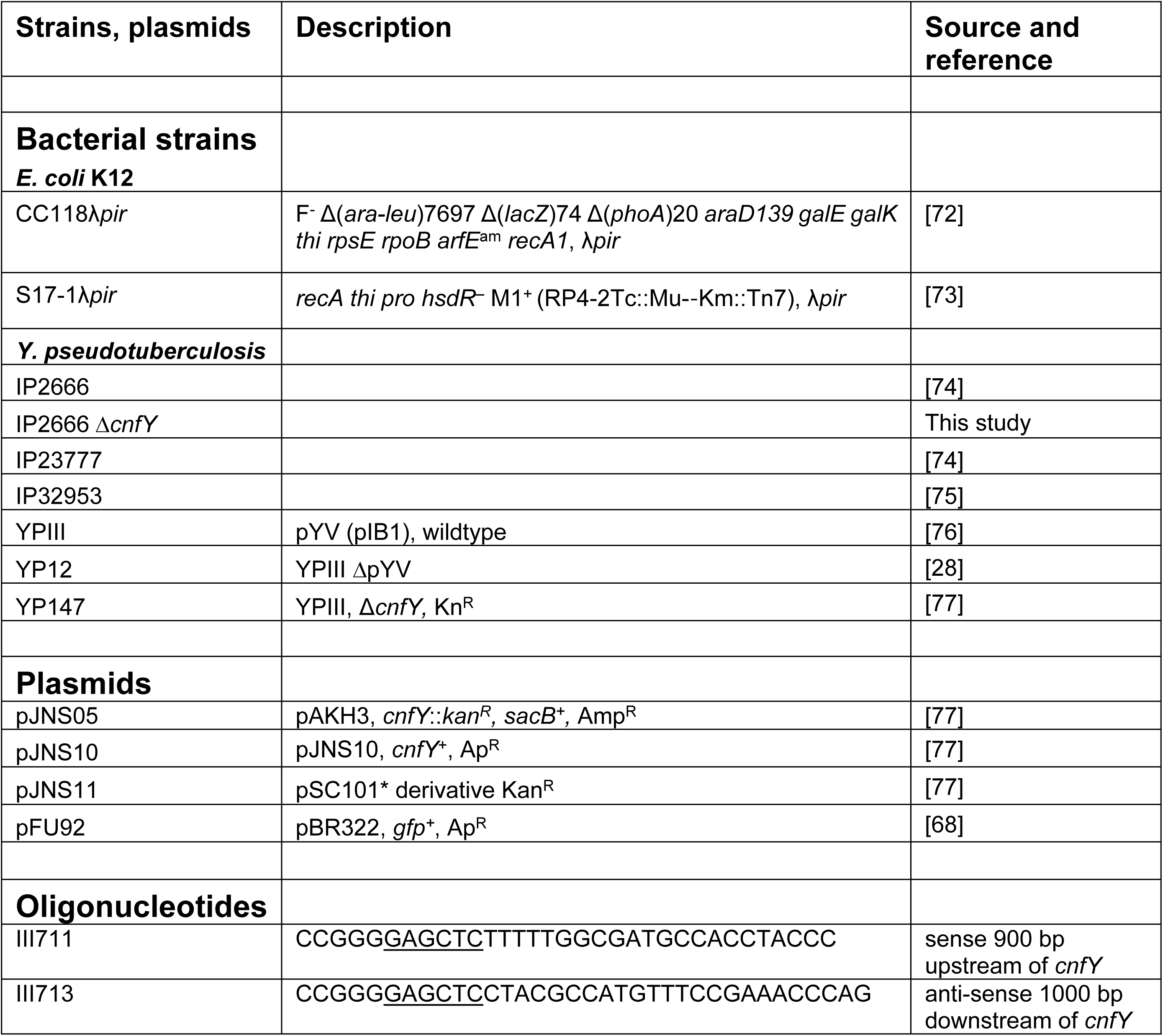
Bacterial strains, plasmids, and oligonucleotides.

## Supplementary Notes

**Note A: Technical details on microscopy and imaging**

For confocal microscopy, an HC PL APO 63x/1,40 Glyc CS2 objective (Leica) was used within a Leica Stellaris 5. Multiple FOVs were recorded in four different channels using unidirectional scanning with a Zoom of 3.0 to capture a sufficient number of exit events. The Area was scanned at a speed of 600 with a resolution of 512 x 512 px. To decrease the signal-to-noise ratio, a frame averaging of 3 was used for each Fluorophore. The pinhole was set to 102.9 μm (1 AU, 580 nm).

The Fluorophore CellMask deep red was excited at 661 nm with an Intensity of 2.00. The Emission was detected within 667 and 839 nm at a Gain of 99.9 and an Offset of 0.

DAPI was excited at 405 nm with an Intensity of 1.48. The Emission was detected within 420 and 495 nm at a Gain of 2.5 and an Offset of 0.

EGFP was excited at 405 nm with an Intensity of 1.48. The Emission was detected within 420 and 495 nm at a Gain of 2.5 and an Offset of 0.

Alexa Fluor 568 was excited at 495 nm with an Intensity of 4.00. The Emission was detected within 504 and 647 nm at a Gain of 225.4 and an Offset of 0.

**Note B: Technical details on adjustment and exportation of confocal images** Images were adjusted using LASX software (Version 4.7.0.28176) to minimize the signal-to-noise ratio and reduce the background. Therefore, the histograms were limited for DAPI within 20 and 255, for EGFP within 10 and 120, for Alexa Fluor 568 within 10 and 160, and for CellMask DeepRed within 20 and 255. The pictures were exported, and the channels were merged with the LASX software as Scaled Vector Graphics. DAPI, EGFP, Alexa Fluor 568, and CellMask DeepRed were merged and exported.

